# Rap1 uses Canoe-dependent and Canoe-independent mechanisms to regulate apical contractility and allow embryonic morphogenesis without tissue disruption

**DOI:** 10.1101/2022.05.18.492492

**Authors:** Kia Z. Perez-Vale, Kristi D. Yow, Melissa Greene, Noah J. Gurley, Mark Peifer

## Abstract

Embryonic morphogenesis is powered by dramatic changes in cell shape and arrangement, driven by the cytoskeleton and its connections to adherens junctions. This requires robust linkage, allowing morphogenesis without disrupting tissue integrity. The small GTPase Rap1 is a key regulator of cell adhesion, controlling both cadherin-mediated and integrin-mediated processes. We have defined multiple roles in morphogenesis for one Rap1 effector, Canoe/Afadin, which ensures robust junction-cytoskeletal linkage. We now ask what mechanisms regulate Canoe and other junction-cytoskeletal linkers during *Drosophila* morphogenesis, defining roles for Rap1 and one of its guanine nucleotide exchange factor (GEF) regulators, Dizzy. Rap1 uses Canoe as one effector, regulating junctional planar polarity. However, Rap1 has additional roles in junctional protein localization and balanced apical constriction—in its absence, Bazooka/Par3 localization is fragmented, and cells next to mitotic cells apically constrict and invaginate, disrupting epidermal integrity. In contrast, the GEF Dizzy has phenotypes similar to but slightly less severe than Canoe loss, suggesting this GEF regulates Rap1 action via Canoe. Taken together, these data reveal that Rap1 is a crucial regulator of morphogenesis, likely acting in parallel via Canoe and other effectors, and that different Rap1 GEFs regulate distinct functions of Rap1.

## Introduction

Small GTPases in the Ras superfamily regulate virtually all aspects of cell biology, ranging from receptor tyrosine kinase signaling to vesicular trafficking to nuclear import to cell adhesion (Cherfils and Zeghouf, 2013). Small GTPases cycle between GTP-bound (ON) states and GDP-bound (OFF) states, with the transitions promoted by guanine nucleotide exchange factors (GEFs), which enhance nucleotide exchange and thus turn GTPases on, and GTPase activating proteins (GAPs) which stimulate GTPase activity to promote deactivation. Active GTPases bind to and activate diverse effector proteins, stimulating downstream events. The small GTPase Rap1 belongs to the Ras superfamily, and acts as a key regulator of cell adhesion, controlling both cadherin-mediated and integrin-mediated processes (Boettner and Van Aelst, 2009). Our recent work defined key roles for the Rap1 effector Canoe (Cno), homolog of mammalian Afadin, in morphogenesis. We now want to move upstream, defining the roles of Rap1 itself, its upstream regulators, and other potential Rap1 effectors implicated in regulating cell shape change and cell migration during embryonic development.

Rap1 is a conserved GTPase that plays important roles across the animal kingdom. The two mammalian Rap1 proteins regulate diverse processes ranging from neuronal migration to leukocyte trafficking to platelet activation. Rap1 regulation of blood vessel development and homeostasis, promoting angiogenesis and endothelial barrier function, provides an illustrative example (Chrzanowska-Wodnicka, 2017). In this tissue, Rap1 acts downstream of VE-cadherin and the receptor tyrosine kinase VEGFR, and upstream of multiple effectors, including Rasip1, Krit-1, and Afadin. In leukocytes and platelets, Rap1 acts via RIAM and Talin to modulate integrin function (Bromberger *et al*., 2018; Lagarrigue *et al*., 2020). Mammals have a large suite of Rap1 GEFs, which act in different tissues and times—for example the GEF EPAC activates Rap1 to ensure endothelial barrier function (Pannekoek *et al*., 2020), while RA-GEF-1 regulates Rap1 to promote callosal axons crossing the midline (Bilasy *et al*., 2011) and RA-GEF-2 regulates Rap1 during spermatogenesis (Okada *et al*., 2014).

*Drosophila* Rap1 was first identified via dominant gain-of-function mutations affecting development of the stereotyped ommatidial cell arrangements in the developing eye (Hariharan *et al*., 1991; Asha *et al*., 1999; Walther *et al*., 2018). Rap1 was subsequently implicated in regulating diverse cellular events in *Drosophila*. Rap1 regulates cell adhesion in the larval wing disc (Knox and Brown, 2002) and the germline stem cell niche (Wang *et al*., 2006), and planar cell polarity and cell shape in the pupal wing (O’keefe *et al*., 2009). It regulates migration of somatic border cells in the ovary (Sawant *et al*., 2018), embryonic macrophages (Huelsmann *et al*., 2006), germline precursors (Asha *et al*., 1999) and embryonic mesodermal cells (Asha *et al*., 1999; McMahon *et al*., 2010). Rap1 also regulates organ development and function, including synaptogenesis at neuromuscular junctions (Ou *et al*., 2019), axon guidance (Yang *et al*., 2016), epidermal muscle attachment (Camp *et al*., 2018), and nephrocyte function (Dlugos *et al*., 2019).

Given these diverse roles of *Drosophila* Rap1, scientists explored which GEFs regulate Rap1 in different contexts. Several GEFs were implicated in different events. C3G mutants have defects in body wall muscle development (Shirinian *et al*., 2010) and nephrocyte function (Dlugos *et al*., 2019), while inappropriate C3G activation alters cell patterning in the eye and wing (Ishimaru *et al*., 1999) The Rap1 GEF EPAC is implicated in Malpighian tubule function (Efetova *et al*., 2013), in the role of mushroom body Kenyon cell neurons in aversive learning (Richlitzki *et al*., 2017), and in the response to anthrax toxin, via a role in blocking fusion of recycling endosomes with the plasma membrane (Guichard *et al*., 2017). While these two GEFs have relatively limited roles, Dizzy (Dzy), also known as PDZ-GEF, has broader roles. Dizzy is the GEF that mediates Rap1’s roles in macrophage (Huelsmann *et al*., 2006) and border cell migration (Sawant *et al*., 2018), germline stem cell adhesion (Wang *et al*., 2006), and synapse development and function at the neuromuscular junction (Heo *et al*., 2017; Ou *et al*., 2019), while in the developing eye and wing Dzy acts in parallel with the atypical GEF Sponge (Lee *et al*., 2002; Eguchi *et al*., 2013).

Our interest is in the regulation of cell adhesion and the cytoskeleton during embryonic morphogenesis, using *Drosophila* as a model. These events provide an outstanding place to explore the complexities of the Rap1 pathway, and define its regulators and effectors (Fig. 1A). We have focused on the *Drosophila* Rap1 effector Cno. Cno plays important roles in linking cell-cell adherens junctions (AJs) to the cytoskeleton, reinforcing cell junctions under mechanical tension. Through this function, Cno regulates initial apical positioning of AJs during polarity establishment (Choi *et al*., 2013; Bonello *et al*., 2018), apical constriction of mesodermal cells (Sawyer *et al*., 2009), cell intercalation during germband elongation (Sawyer *et al*., 2011; Perez-Vale *et al*., 2021), dorsal closure (Boettner *et al*., 2003; Choi *et al*., 2011; Choi *et al*., 2013), and epithelial integrity (Sawyer *et al*., 2009; Manning *et al*., 2019). However, we and others have only just begun to explore the roles of Rap1 and its GEFs in these events, to define in which events Rap1 uses Cno as its effector, and to determine which GEFs activate Rap1 in these contexts. Thus far, these efforts focused almost exclusively on Cno’s initial roles. Examination of *Rap1* maternal/zygotic mutants revealed Rap1uses Cno as an effector during polarity establishment, mesoderm invagination, and potentially dorsal fold formation (Asha *et al*., 1999; Spahn *et al*., 2012; Choi *et al*., 2013; Wang *et al*., 2013; Bonello *et al*., 2018). The GEF Dzy plays an important role in these events (Spahn *et al*., 2012; Bonello *et al*., 2018). However, things are complex, as the GEF Sponge is also involved in apical-basal positioning of AJs (Schmidt *et al*., 2018), and *dzy* mutants only mediate a subset of the effects of Rap1 mutants during polarity establishment (Bonello *et al*., 2018). Use of dominant negative and activated variants also support the idea that Rap1 acts via Cno during collective cell migration during dorsal closure (Boettner *et al*., 2003), and there has been some examination of the effect *dzy* zygotic mutants on dorsal closure (Boettner and Van Aelst, 2007), but beyond this we do not know what roles Rap1 or Dzy play in many events of morphogenesis that require Cno, including cell intercalation during germband extension and maintenance of epidermal integrity. To fully define the role of Rap1 signaling during these key processes, we need to address two key knowledge gaps: Is Cno the sole Rap1 effector during morphogenesis, or does Rap1 have broader effects through other effectors (Fig. 1A)? Is Dzy the relevant Rap1 GEF during these stages, or do other GEFs play roles (Fig. 1A)? To address these knowledge gaps, we set out to define the roles of Rap1 and Dizzy after gastrulation, interrogating which events occur via a Rap1-Cno pathway, what roles Rap1 plays that require other effectors, and in which Cno-regulated events Dzy is the GEF involved in Cno activation.

**Figure 1.**
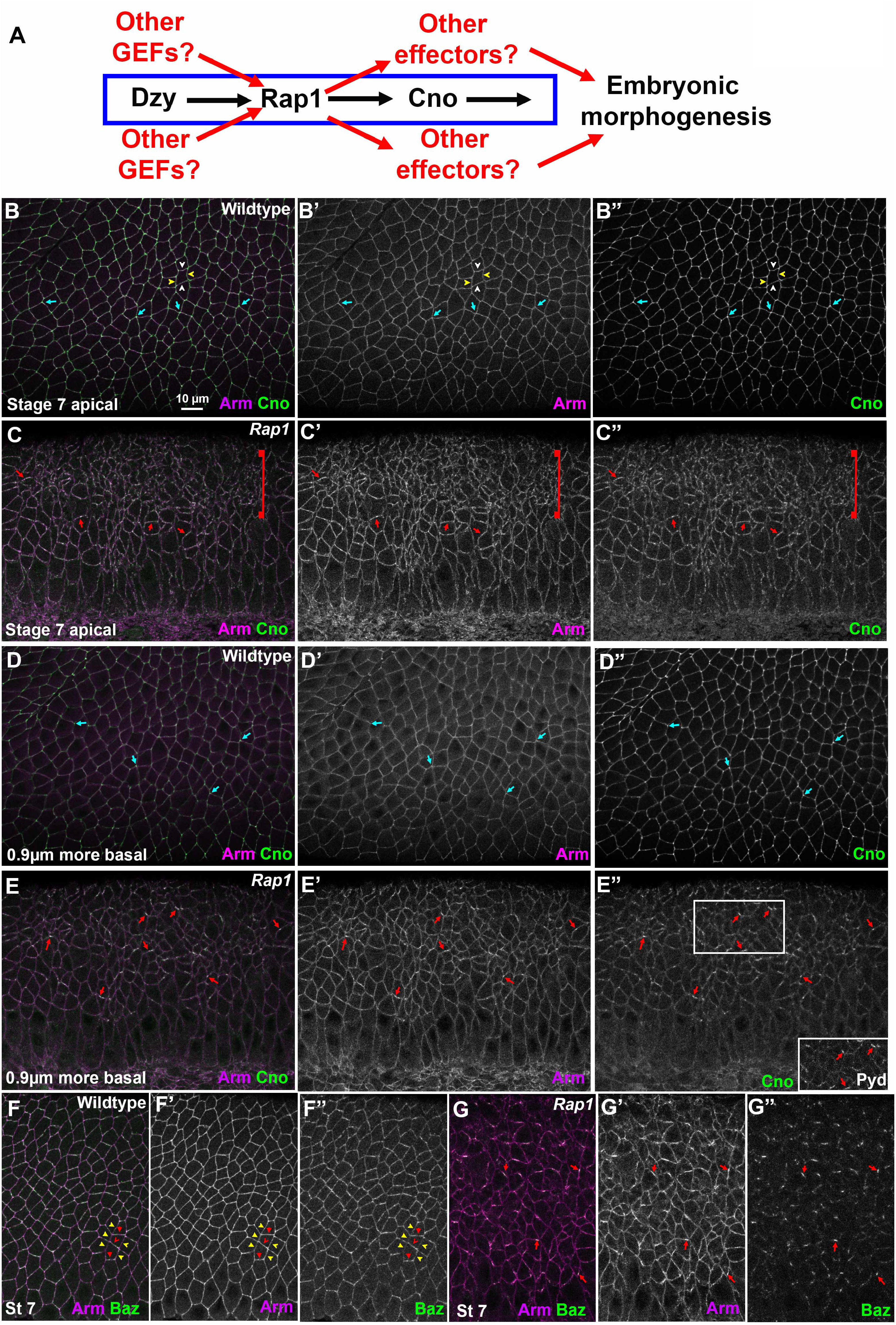
*Rap1* RNAi rapidly leads to alterations in apical junctions, variability in apical cell area, and fragmentation of junctional Baz. A. Diagram of the Rap1 “pathway”. B-G. Stage 7 embryos, imaged at the level of apical AJs (B, C, F, G) or 0.9µm more basal (D, E). Unless noted, in this Figure and all others embryos are anterior left and dorsal up. B, D. Wildtype. Apical cell area is relatively similar at both sectional planes. Cno is enriched at TCJs (cyan arrows). C. *Rap1* RNAi. Many cell junctions become difficult to follow in the apicalmost plane (brackets), and at other AJs Arm and Cno staining become more punctate (red arrows). E. *Rap1* RNAi. More basally cell areas remain much more variable than in wildtype, and junctional accumulation of Arm, Cno and Pyd is not as continuous as in wildtype (red arrows). F. Wildtype. Baz is found all around the cell circumference, but is planar polarized to DV borders (red arrowheads) as opposed to AP borders (yellow arrowheads). G. *Rap1* RNAi. Baz staining is fragmented, and regions where it remains also accumulate elevated levels of Arm (arrows).

## Results

### While *Rap1* RNAi mimics the effect of Cno loss on AJ planar polarity, it also dramatically alters apical AJs and destabilizes Bazooka/Par3 localization

Our goal was to explore the roles of Rap1 and its regulators during morphogenesis and contrast these with the roles of the Rap1 effector Cno. Previous analysis of the role of Rap1 in embryos was largely confined to the earliest events of morphogenesis—initial establishment of apical-basal polarity and invagination of the mesoderm, the first event of gastrulation (Sawyer *et al*., 2009; Spahn *et al*., 2012; Choi *et al*., 2013). In these events, Rap1 and Cno loss lead to similar defects, but since Cno is only one of Rap1’s effectors, we suspected Rap1 might use other effectors to carry out additional roles in junctional integrity as AJs are challenged by cell intercalation, cell division and neuroblast invagination (Fig. 1A). Cuticle analysis was potentially consistent with this hypothesis, as the Rap1 cuticle phenotype is as or more severe than that of most *cno* mutants (Bonello *et al*., 2018). We thus examined the effects of Rap1 loss, using a previously validated RNAi line that, when driven by the *matGAL4* driver, reduces Rap1 protein to undetectable levels through the end of dorsal closure (Bonello *et al*., 2018). The effects of *Rap1* RNAi on apical-basal polarity establishment match those of *Rap1* maternal/zygotic mutants (Choi *et al*., 2013; Bonello *et al*., 2018), and *Rap1* RNAi leads to fully penetrant defects in mesoderm invagination (31/31 stage 7 and 8 embryos), another previously characterized *Rap1* phenotype (Spahn *et al*., 2012). *Rap1* RNAi thus is a well validated reagent to substantially reduce or eliminate Rap1 function.

To determine whether Rap1 primarily acts via Cno or has additional effectors during embryonic morphogenesis, we compared effects of *Rap1* RNAi with those of Cno loss. We first examined germband extension, an embryonic event in which cells in the ectoderm undergo cell intercalation, which in turn helps narrow and extend the germband (Kong *et al*., 2017). This process is driven by reciprocal planar polarization of actin and myosin to anterior/posterior (AP) cell borders and AJ proteins to dorsal/ventral (DV) cell borders. Myosin drives constriction of AP borders, creating T1 transition or rosettes, which then rearrange to complete intercalation. Cno is enriched at the sites where AJs are under elevated molecular tension—AP borders and TCJs (Sawyer *et al*., 2009; Sawyer *et al*., 2011; Yu and Zallen, 2020). Cno is important to reinforce these AJs, and in Cno’s absence junctions separate at these sites (Sawyer *et al*., 2011). The fly ZO-1 homolog Polychaetoid (Pyd) is enriched at DV borders with AJ proteins and acts in parallel with Cno to maintain epithelial integrity (Manning *et al*., 2019).

We began our exploration of the mechanisms by which Rap1 regulates morphogenesis, by examining whether it regulates AJ protein localization and stability. In wildtype embryos, the fly beta-catenin homolog Armadillo (Arm) and Cno rapidly transition from the spot AJs seen during cellularization to a more uniform localization along apical junctions as germband extension accelerates during stage 7 (Fig. 1B). Cno is enriched at TCJs (Fig. 1B, cyan arrows) and at AP borders (Fig. 1B, yellow vs white arrowheads; quantified in Fig. 2C), while Arm and Pyd are more uniformly localized between AP and horizonal borders (Fig. 1B; quantified in Fig. 2A, B). Bazooka (fly Par3; Baz) is obviously planar polarized at this stage (Fig. 1F, quantified in Fig. 2D), but remains present on both AP and DV cell borders. In contrast, Cno loss enhances the planar polarization of Arm, Pyd and especially Baz, by strongly reducing their accumulation on AP borders. This weakens AJ-cytoskeletal connections there, leading to apical gaps at AP borders and TCJ (Sawyer *et al*., 2011; Manning *et al*., 2019). However, core AJ proteins remain localized to AJs in *cno* mutants.

**Figure 2.**
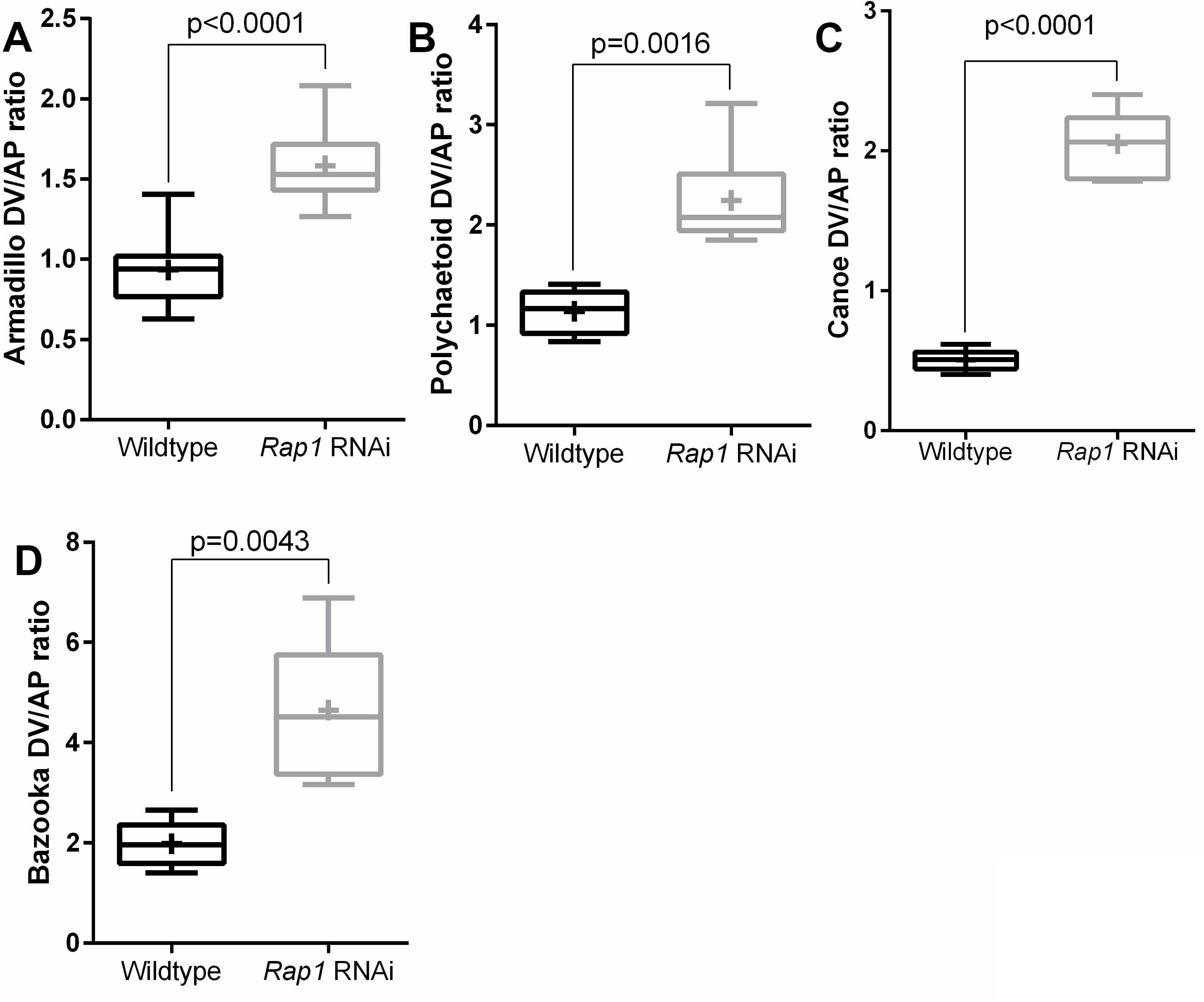
In cells where shape is not drastically altered, *Rap1* RNAi alters junctional protein planar polarity in ways similar to loss of Cno. A-D. Planar polarity quantification. *Rap1* RNAi enhances planar polarity of Arm, Pyd and Baz, and flips the planar polarity of Cno. Numbers of cells analyzed are in Table S1.

AJ protein localization is altered early in Rap1 mutants. While Arm and Cno return to apical AJs in *Rap1* RNAi embryos as gastrulation begins (Bonello *et al*., 2018), we found that their junctional localization was abnormal. This was most striking apically, where in *Rap1* RNAi embryos both Arm and Cno localization to cell borders became more punctate (Fig. 1C). Their localization was less continuous along less affected cell junctions (Fig. 1C, arrows), while their localization to other cell junctions was sufficiently disrupted to make cell borders hard to define at the most apical end of many cells (Fig. 1C, bracket; observed in 16/16 stage 7 embryos). Cell shapes were more regular 0.9µm basally, but AJ junction localization of Arm was less continuous than in the wildtype, and this was even more striking for Cno (Fig. 1D vs E, arrows). Sites where junctional Cno was more intense were also places where both Arm and Pyd were also enriched (Fig. 1E”, inset). The effect of *Rap1* RNAi on Baz localization was even more dramatic. Baz localization already was highly fragmented at stage 7 in *Rap1* RNAi embryos, with just small regions of AJs retaining Baz (Fig. 1G, arrows)— again, these often coincided with the regions where Arm localization was most intense. The strong disruption of apical junctions in many cells and the dramatic fragmentation of Baz were all phenotypes that were absent in *cno* null mutants (Sawyer *et al*., 2011; Manning *et al*., 2019).

We next measured the effect of *Rap1* RNAi on AJ protein planar polarity, choosing the subset of stage 8 cells that were less affected by the dramatic cell shape changes described above. *Rap1* RNAi mimicked the effect of loss of Cno on Arm and Pyd localization (Sawyer *et al*., 2011; Manning *et al*., 2019), elevating planar polarization of Arm (Fig. 2A) and Pyd (Fig. 2B) to DV borders. *Rap1* RNAi also reversed the planar polarity of Cno (Fig. 2C). In each case these alterations reflected the fact that protein localization to AP borders was reduced. In the subset of cells where we could measure Baz, its planar polarity was also strongly enhanced (Fig. 2D). Together these data are consistent with Cno being one of Rap1’s effectors during these stages, as their effects on junctional planar polarity were similar in direction. However, the dramatically more severe effects of Rap1 loss on apical cell junctions and on Baz junctional integrity strongly suggest Cno is not its sole effector.

### *Rap1* RNAi leads to major defects in balanced apical contractility and cell shape regulation

We next explored more broadly the effects of *Rap1* RNAi in cell shape change and AJ integrity as morphogenesis continued. Until the onset of germband extension, *Rap1* RNAi embryos were virtually normal (Fig. 3A, B), like *Rap1* mutants, with only modest alterations in the regularity of apical cell area during cellularization (Fig. 3A, arrows; Choi *et al*., 2013). However, as germband extension accelerated during stage 7, *Rap1* RNAi embryos began to exhibit striking defects. During stage 7, all cells assemble both a junctional and an apical contractile actomyosin cytoskeleton. In wildtype, contractility is relatively balanced. While some cell stretching is seen in the wildtype next to the invaginated mesoderm, cell shapes remain largely regular (Fig. 3C, E), with relatively consistent apical areas even as individual cell borders contract to mediate cell intercalation. In contrast, in *Rap1* RNAi embryos apical cell shapes became highly abnormal (Fig. 3C vs D). In wildtype, cell shapes are consistent, from the apical ends of the cells to more basal slices (Fig. 3E-E”). In *Rap1* RNAi embryos the alteration in cell shape was most striking in the apical-most region of the cells, where cell borders became difficult to define (Fig. 3F-F”). Cell borders were more clearly visible just 0.6µm more basal in *Rap1* RNAi embryos, but cell cross-sectional areas were highly variable, with apically constricted cells (Fig. 3F, F’, red arrows) adjacent to cells with much larger apical areas (Fig. 3F, F’ cyan arrows). In a subset of embryos, apically constricted cells were aligned along the anterior-posterior axis, leading to groups of cells beginning to fold inward (Fig. 3D, yellow arrow; 9/12 stage 7 embryos had at least one such fold). In contrast, while *cno* null mutants do exhibit abnormal cell shapes and AJ gaps at AP borders and TCJs (Fig. 3G; (Sawyer *et al*., 2011), they do not share the extreme alterations in apicalmost AJs seen after *Rap1* RNAi.

**Figure 3.**
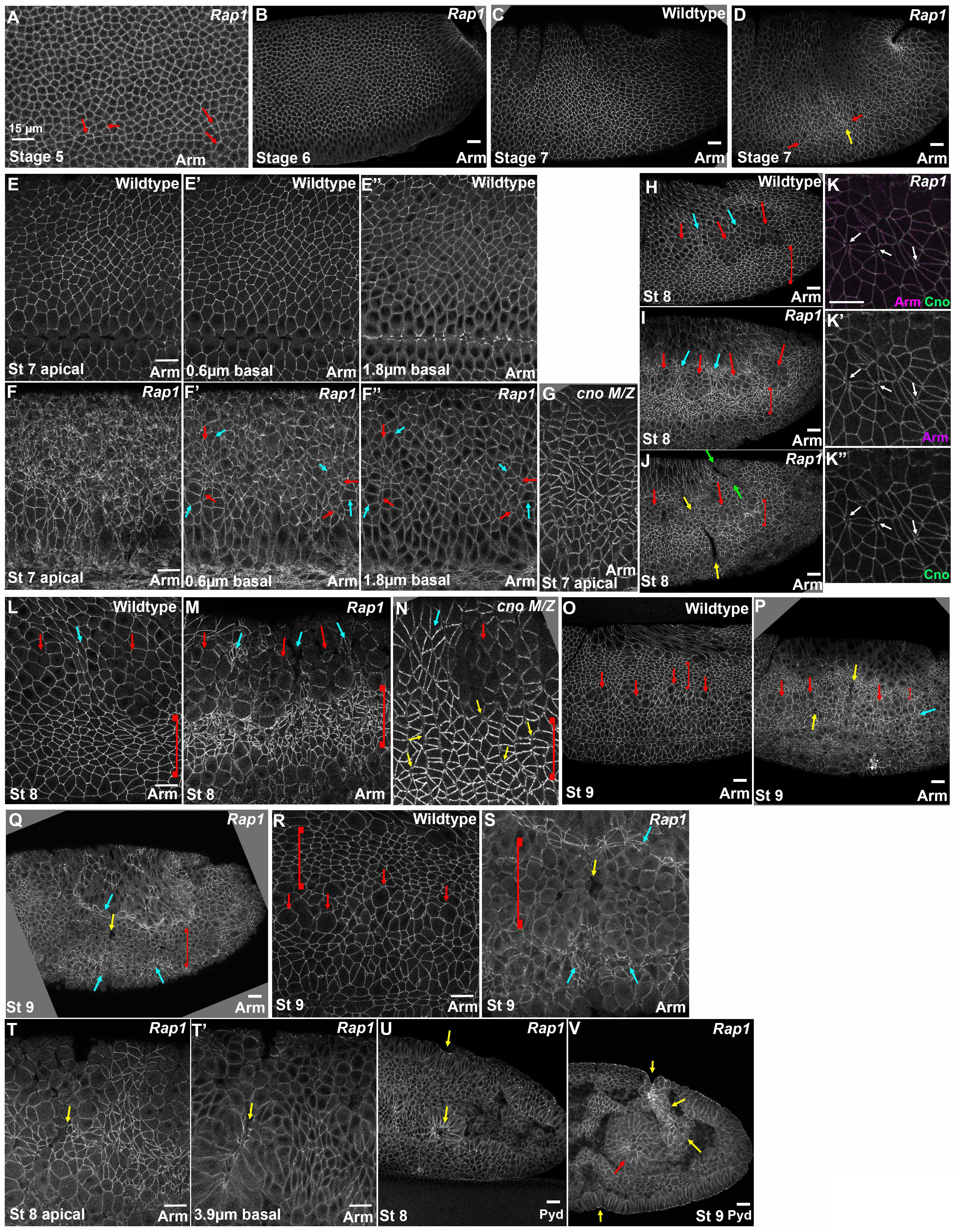
After *Rap1* RNAi apical cell contractility becomes progressively unbalanced. A, B. At stages 5 and 6, apical cell areas in *Rap1* RNAi embryos remain relatively uniform with only modest variation. C, D. At stage 7, while apical area remains relatively uniform in wildtype (C), in *Rap1* RNAi embryos groups of cells begin to apically constrict relative to their neighbors (D, arrows). E-G. Stage 7 closeups. In the most apical sections, junctions are difficult to trace in *Rap1* RNAi embryos (E vs F). More basally, junctions are more apparent but apical cell areas are quite variable relative to wildtype, with small (F’, F” red arrows) and large (cyan arrows) cross sectional areas. G. *cno M/*Z null mutants do not exhibit the dramatic defects in AJs in the apical plane. H-N. Stage 8. In wildtype, cells round up to divide in mitotic domain 11 (H, L red arrows), and, cells between or adjacent undergo some apical constriction (H, L cyan arrows and brackets). K. *Rap1* RNAi. In less affected regions gaps in AJs open at TCJs and AP borders (arrows). I, J, M. In *Rap1* RNAi embryos, cells between and adjacent to mitotic domains are hyper-constricted (I, J, M cyan arrows and brackets) and at times constriction leads to deep folds (J, yellow and green arrows). N. *cno M/*Z null mutants have gaps between cells at AJs under tension (yellow arrows), but do not exhibit the dramatic accentuation of apical constriction of cells ventral to mitotic domain 11 (bracket). O-S. Stage 9. In wildtype (O, R), dorsal ectodermal cells that divided in mitotic domain 11 resume columnar architecture (brackets) while cells in mitotic domain N round up for division (arrows). In milder *Rap1* RNAi embryos (P), both dorsal ectodermal cells (bracket) and cells ventral to the mitotic cells (cyan arrow) are hyper-constricted, and folds remain (yellow arrows). Other *Rap1* RNAi embryos have more severe defects. (Q, S). Many mitotic domain 11 and domain N cells remain rounded up (bracket) and flanking cells are very hyper-constricted (cyan arrows) and are folding inward (yellow arrows). T-V. Examples of deep epithelial folds after *Rap1* RNAi (yellow arrows) descending to the level of the posterior midgut (red arrow).

The cell shape defects after *Rap1* RNAi became even more pronounced in stage 8. *Drosophila* cells divide in programmed groups called mitotic domains. During embryonic stage 8in wildtype, mitotic domain 11 becomes prominent in the thoracic and abdominal regions and cells in this mitotic domain round up to divide in (Fig. 3H, L, red arrows). In the wildtype, cells between (Fig. 3H, L, cyan arrows) or more ventral to the mitotic domains (Fig. 3H, L, brackets) apically constrict slightly, presumably due to reduced pulling forces from mitotic neighbors. In *Rap1* RNAi embryos these differences were strongly exaggerated, with cells between (Fig. 3I, M, cyan arrows) or more ventral to the mitotic domains (Fig. 3I, M brackets) becoming irregular in shape and highly apically constricted. This phenotype was fully penetrant (17/17 stage 8 embryos). The folds observed during stage 7 also became more prevalent and deeper, with folds extending across the ectoderm (Fig. 3J, yellow arrows) and sometimes even extending into the amnioserosa (Fig. 3J, green arrows; 17/18 stage 8 embryos had folds). In less affected regions of the ectoderm, gaps formed at TCJs and AP cell borders (Fig. 3K), similar to those seen in *cno* mutants (Sawyer *et al*., 2011; Perez-Vale *et al*., 2021). While *cno* null mutants continue to exhibit altered cells shapes and gaps in apical AJs during stage 8 (Fig. 3N; Sawyer *et al*., 2011; Manning *et al*., 2019), they lack the dramatically imbalanced apical contractility seen after *Rap1* RNAi.

In wildtype embryos during stage 9, dorsal ectodermal cells in mitotic domain 11 resume columnar shape (Fig. 3O, R, brackets) while cells in mitotic domain N begin to round up to divide (Fig. 3O, R, arrows). In *Rap1* RNAi embryos epithelial defects continued or worsened at this stage. In less severe embryos (14/23 scored), dorsal ectodermal cells were hyper-constricted (Fig. 3O vs P, brackets) and grooves remained (Fig. 3P, yellow arrows). In more severe embryos (9/23 scored), dorsal ectodermal cells failed to resume columnar architecture after their division at stage 8 (Fig. 3Q, R vs S, brackets). Adjacent cells became even more apically constricted (Fig. 3Q, S, cyan arrows) or folded into the embryo (Fig. 3Q, S, yellow arrows). These infolding cells formed epithelial balls or folded sheets (Fig. 3T, T’, U arrows), which could become quite extensive (Fig. 3V, yellow arrows), matching the posterior gut in depth and extent (Fig. 3V, red arrow). Once again, this deep infolding seen after In *Rap1* RNAi was not observed after Cno loss (Manning *et al*., 2019).

The alterations in AJ protein localization observed at stages 7 and 8 continued during stage 9. At this stage in wildtype, dorsal ectodermal cells have regained columnar architecture after having divided (Fig. 4A, red bracket), cells in mitotic domain N are rounding up to divide (Fig. 4A, cyan bracket), while the most ventral cells have yet to divide (Fig. 4A, yellow bracket). In *Rap1* RNAi embryos, virtually all dorsal cells remained rounded up, with Arm, Cno and Pyd reduced at AJs in those cells (Fig. 4B, red and cyan brackets). In the more ventral, highly apically constricted cells, Arm, Cno and Pyd localized less uniformly to AJs (Fig. 4B, yellow bracket). Junctional Baz remained very fragmented, even in those cells that retained more columnar architecture (Fig. 4C vs D). Thus, Rap1 is essential for maintaining uniform localization of Arm, Cno and Pyd along AJs, and for continued Baz localization to AJs.

**Figure 4.**
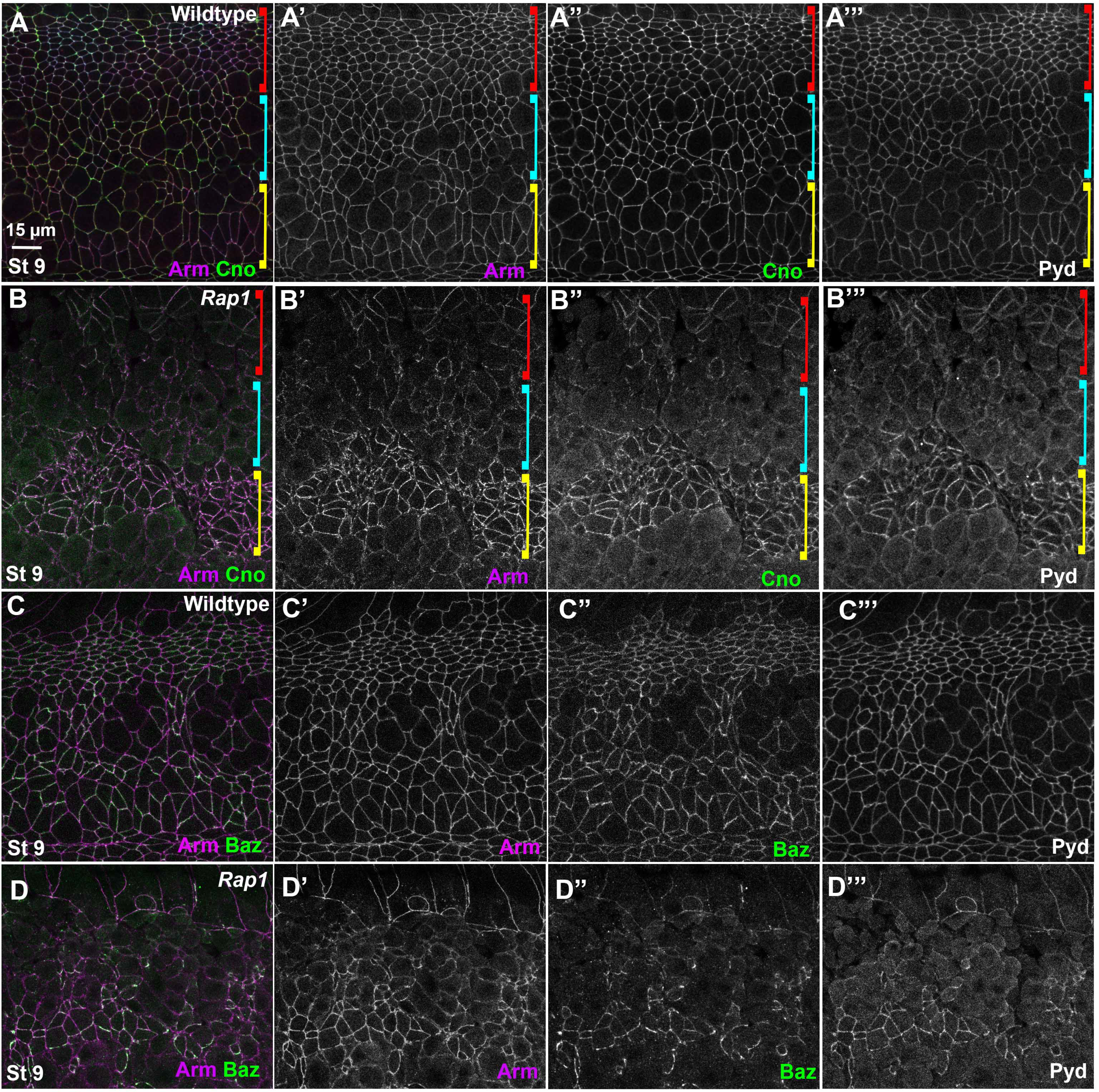
At stages 8 and 9, while core AJ proteins remain at cell junctions, defects in balanced contractility and junctional Baz localization continue or intensify. A,B. In wildtype (A) Arm, Cno, and Pyd remain at junctions in both mitotic and columnar cells. After *Rap1* RNAi (B) junctional Arm, Cno, and Pyd become less continuous in both cell populations. C,D. In wildtype (C) Baz is strong in non-mitotic cells and weaker in mitotic cells. After *Rap1* RNAi (D), Baz localization to junctions is reduced and fragmented.

### Rap1 RNAi leads to major disruption of epidermal integrity that is more severe than that after Cno loss

These cell shape and junctional defects after *Rap1* RNAi ultimately had striking effects on epidermal integrity. By stage 11, phenotypes diverged somewhat, with more and less severely affected embryos, likely due to differences in the zygotic copy number of the RNAi construct (0-2 copies). However, all *Rap1* RNAi embryos had defects in epithelial integrity. At this stage, the wildtype epidermis is continuous from the amnioserosa dorsally to the ventral midline (Fig. 5A). In the more severely affected *Rap1* RNAi embryos epidermal integrity was largely lost by stage 11, with the ventral and lateral epidermis completely disrupted, and only patches of dorsal epidermis remaining intact (Fig. 5B, green arrows). Laterally and ventrally one could see the apical ends of cells that have folded into the interior (Fig. 5B, red arrows). Less severely affected embryos had similar defects, but the amount of remaining dorsal epidermis was more extensive (Fig. 5C). *cno M/Z* null mutant embryos also have defects in the ventral epidermis, leading to eventual holes in the ventral cuticle, but the amount of remaining epidermis is substantially more extensive (Fig. 5D; Sawyer *et al*., 2009; Sawyer *et al*., 2011; Manning *et al*., 2019; Perez-Vale *et al*., 2021). After stage 11, many *Rap1* RNAi embryos had such substantial disruptions of the epidermis that they were difficult to stage precisely, because only patches of epidermis remained intact (Fig. 5E, green arrows). In contrast, while *cno M/Z* null mutant embryos have defects in head involution, dorsal closure and ventral epidermal integrity at stage 13-14, their lateral and dorsal epidermis remains intact (Fig. 5F; Sawyer *et al*., 2009; Sawyer *et al*., 2011; Manning *et al*., 2019; Perez-Vale *et al*., 2021). Thus, Rap1 is essential for maintaining epithelial integrity during morphogenesis.

**Figure 5.**
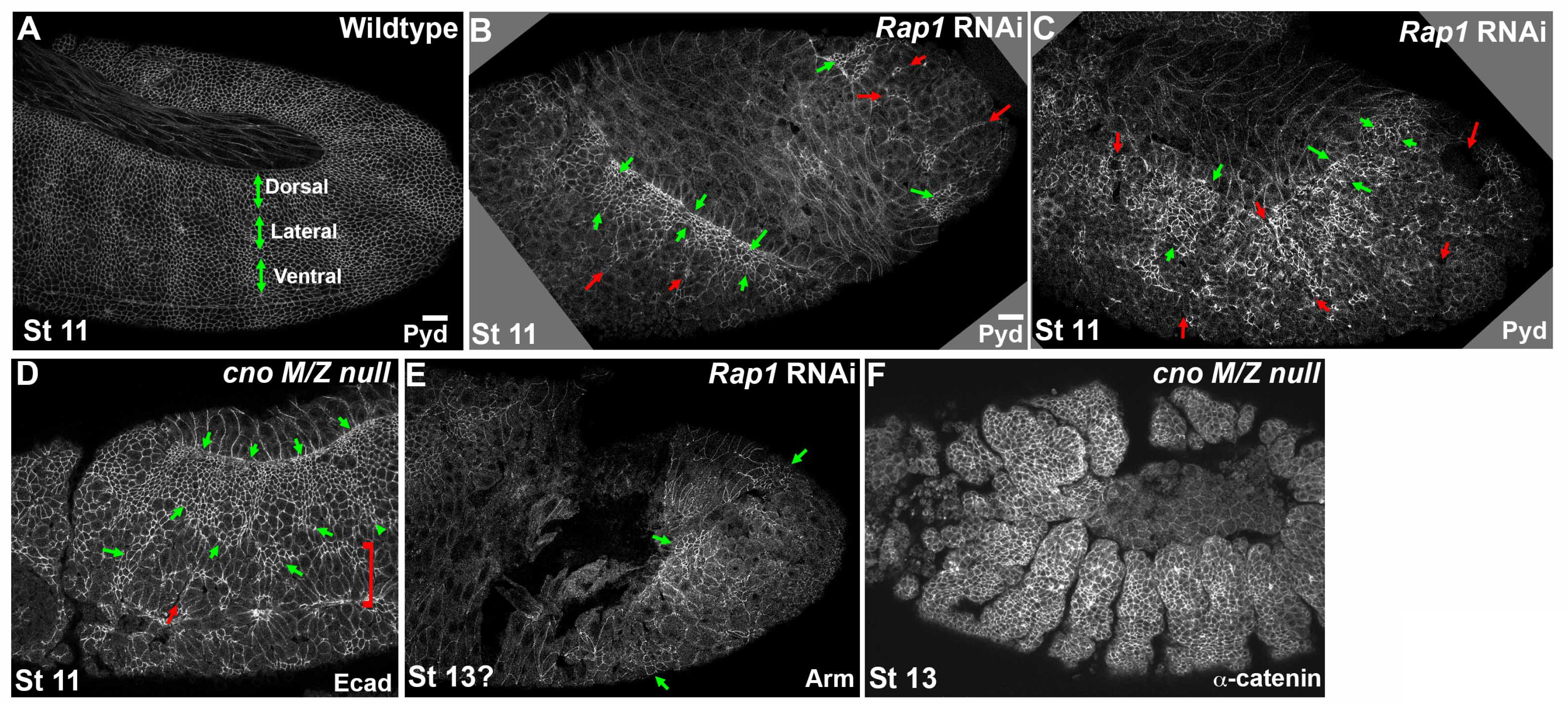
*Rap1* RNAi leads to widespread disruption of epidermal integrity, more severe than that seen after Cno loss. A-D. Stage 11. A. The wildtype epidermis remains intact despite segmental cell shape change and tracheal pit invagination. Dorsal, lateral and ventral epidermal regions are indicated. B. In severe *Rap1* RNAi embryos only small regions of intact dorsal epidermis remain near the amnioserosa (green arrows). The lateral and ventral epidermis are totally disrupted, with the apical ends of infolded cells seen in some places (red arrows). C. Similar though somewhat less drastic defects are seen in other *Rap1* RNAi embryos. D. In *cno M/Z* null mutants the ventral most epidermis is disrupted (bracket) and occasional infolding is seen (red arrow) but the region of remaining intact epidermis is more substantial (green arrows). E. Later stage *Rap1* RNAi embryo. Staging becomes difficult as only small regions of intact epidermis remain (green arrows). F. In contrast, stage 13 *cno M/Z* null mutants retain intact dorsal and lateral epidermis, but do fail during dorsal closure and have the deep segmental groove phenotype.

Putting all of these analyses together, Rap1 and Cno loss share some features, including parallel defects in junctional protein planar polarity, consistent with Cno being one of Rap1’s effectors. However, the multiple defects seen after *Rap1* RNAi that are not shared by *cno* mutants, including dramatic destabilization of Baz localization, very unbalanced apical contractility, and extreme loss of epidermal integrity, suggest that Rap1 relies on additional effectors during this phase of morphogenesis.

### Developing and validating tools to strongly reduce Dzy function to define its role in morphogenesis

We also want to know which GEFs activate Rap1 during morphogenesis. Dzy is one of several Rap1 GEFs. During cellularization, Dzy loss mimics the effects of loss of Cno on junctional polarization (Bonello *et al*., 2018), and both Cno and Dzy are required for effective apical constriction during mesoderm invagination (Spahn *et al*., 2012), the first event of gastrulation. Thus, Dzy is an important regulator of Rap1/Cno during these early stages. However, Dzy’s roles in other events of embryonic morphogenesis remain essentially unexplored.

We and others previously used the FLP/FRT/DFS technique to generate maternal/zygotic *dzy* mutants (Boettner and Van Aelst, 2007; Spahn *et al*., 2012; Bonello *et al*., 2018), but females lay few eggs, suggesting possible roles in oogenesis. To circumvent this, we sought to create an effective shRNA reagent that we could drive maternally and thus use to deplete both maternal and zygotic Dzy (Staller *et al*., 2013). We and others have used this approach very effectively, in our case to reduce maternal and zygotic expression of Rap1 and Cno (Bonello *et al*., 2018). We generated such an shRNA reagent, using the pWalium22 vector to create an shRNA targeting *dzy’s* 5th exon (the oligos used to target *dzy* are listed in the Methods). We inserted the shRNA plasmid into the right arm of the 3^rd^ chromosome, using phiC31/attP2 site integration. We drove this UAS-*dzy* shRNA construct in the female germline with a strong two-component maternal germline driver line, *matGAL4* (Staller *et al*., 2013). Using this GAL4 driver to drive shRNAs has worked well for us in the past to greatly reduce or eliminate maternal contribution of both Cno and Rap1 (Bonello *et al*., 2018).

Because we could not find a Dzy antibody that worked in immunoblotting, we tested the effectiveness of depleting maternal Dzy by comparing the effects of *dzy* RNAi on cellularization and mesoderm invagination with those previously characterized for maternal/zygotic *dzy* null mutants. We first compared the effects of *dzy* RNAi on AJ polarization during cellularization, a process likely driven exclusively by maternal contribution, comparing it to our previous analysis of *dzy* null maternal mutants (Bonello *et al*., 2018). During cellularization, Cno localizes to nascent apical spot AJs (SAJs; Fig. 6A yellow arrow, C). At the apical position of SAJs, Cno localizes to bicellular junctions and is also enriched at tricellular junctions (TCJs; Fig. 6C, yellow vs cyan arrows). At TCJs strong Cno localization extends more basally (Fig. 6E, yellow vs cyan arrows), forming cable-like structures visible in maximum intensity projections (MIPs; Fig. 6G). While loss of Rap1 leads to complete loss of Cno from the plasma membrane (Sawyer *et al*., 2009), *dzy* null maternal mutants have less severe effects, suggesting that additional Rap1 GEFs may act at this stage (Bonello *et al*., 2018). In *dzy* maternal mutants, while Cno remains enriched in spot AJs and roughly localized to the apicolateral region, its localization spreads more apically and Cno enrichment at TCJs and formation of cable-like structures is lost (Bonello *et al*., 2018). *dzy* RNAi replicated the effect of *dzy* null maternal mutants on Cno localization during cellularization. While Cno remained roughly localized apically (Fig. 6B) and still accumulated in bicellular SAJs (Fig. 6D, F, yellow arrows), Cno localization expanded more apically (Fig. 6B, H, cyan arrows) and enrichment at TCJs was lost, both at the levels of normal nascent spot AJs (Fig. 6D, yellow vs cyan arrows) and 1.5µm basally (Fig. 6F, yellow vs cyan arrows). As a result, in maximum intensity projections, Cno cables were fragmented (Fig. 6H). Thus, *dzy* RNAi replicates all the phenotypes of *dzy* null maternal mutants (Bonello *et al*., 2018), consistent with our RNAi effectively knocking down maternal Dzy. As we had observed with *dzy* null maternal mutants (Bonello *et al*., 2018), apical junctional enrichment of Cno was largely restored as gastrulation began in stage 6 (Fig. 6I vs J). As a second test of the effectiveness of maternal Dzy depletion by *dzy* RNAi, we assessed its effects on mesoderm invagination. In *dzy* null maternal mutants, mesoderm invagination fails (Spahn *et al*., 2012). *dzy* RNAi also had an extremely penetrant effect on this process, with partial (Fig. 6Q vs R, white arrows) or complete (Fig. 6S, white arrows) failure of mesoderm invagination in 52 of 53 embryos (as assessed at stage 7). Together these data suggest that our shRNA is effective at very strongly reducing or eliminating Dzy maternal contribution.

**Figure 6.**
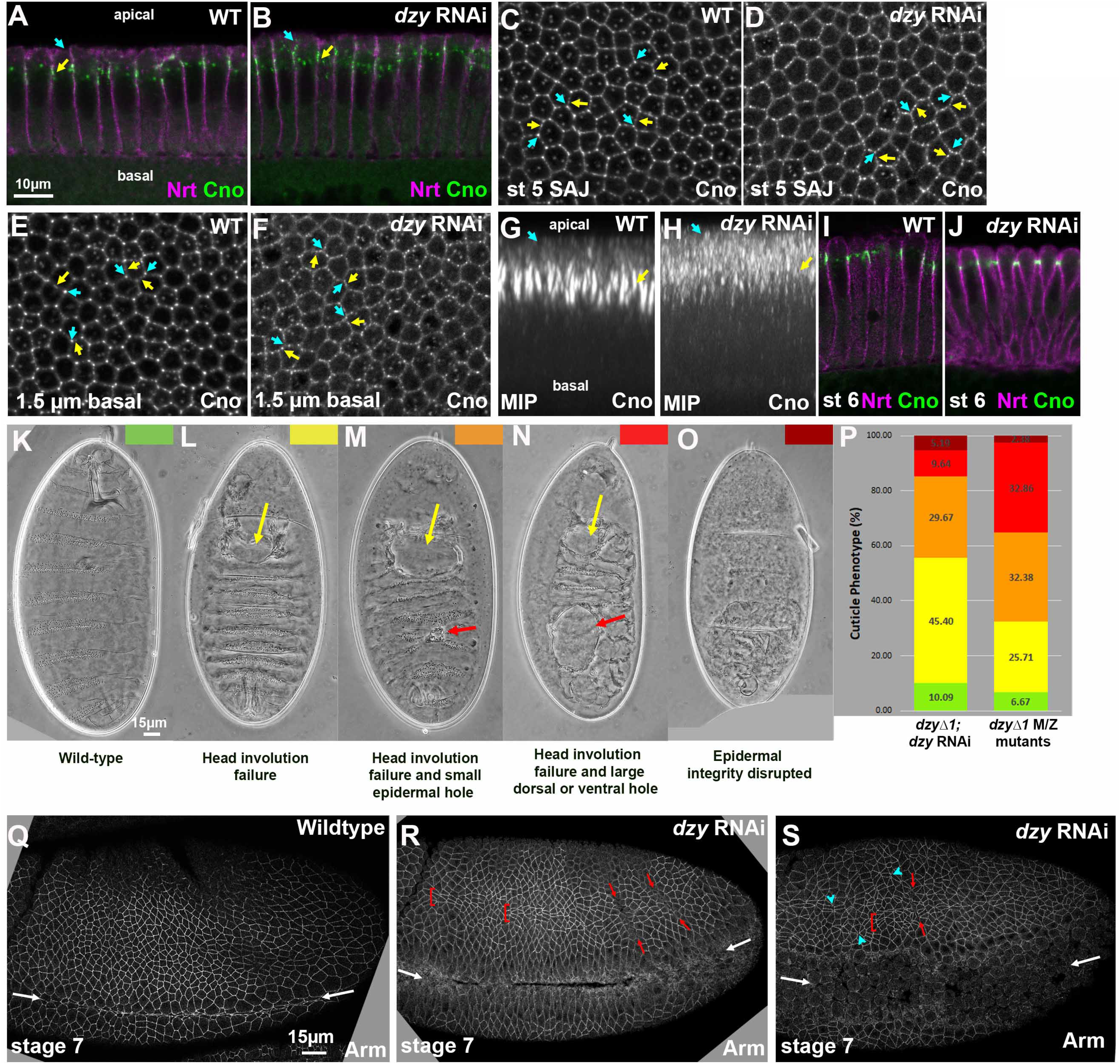
*dzy* RNAi largely mimics total maternal/zygotic loss of Dzy. A, B. Apical-basal cross sections. Neurotactin (Nrt) marks the plasma membrane. Cno remains apically enriched at nascent SAJs after *dzy* RNAi (yellow arrow) but some Cno moves more apically (cyan arrow). C-F. *en face* sections at the level of the SAJs (C, D) or 1.5µm more basal (E, F). In wildtype Cno localizes to all SAJs, but is somewhat enriched at TCJs relative to bicellular junctions (C, cyan vs yellow arrows), and is strongly enriched at TCJs 1.5µm basal (E). *dzy* RNAi leads to loss of TCJ enrichment at both the SAJ level (D) and more basally (F). G, H. Maximum intensity projections of apical-basal cross sections. In wildtype TCJ Cno forms cables (G, yellow arrows) and is largely excluded apically (G, cyan arrows). *dzy* RNAi (H) disrupts TCJ cables and increases levels apical to the SAJs. I, J. Apical-basal cross sections. At gastrulation onset (stage 6) Cno localization returns to its normal apical position after *dzy* RNAi. K-O. Representative embryonic cuticles illustrating different phenotypic categories. P. Cuticle defects in embryonic progeny from the cross in Figure S1 (*dzy*Δ*1; dzy* RNAi) versus those in maternal/zygotic *dzy*Δ*1* mutants. Q-S. Embryos, stages 7. Q. Wildtype stage 7. Mesoderm invagination is complete (white arrows). R, S. *dzy* RNAi. Mild (R) or strong (S) defects in mesoderm invagination are seen in virtually all embryos (white arrows). Some ectodermal cells are abnormally elongated along the AP axis (brackets), ectopic grooves appear (red arrows), and gaps appear at TCJs and AP borders (cyan arrowheads).

To further ensure reduction of both maternal <u>and</u> zygotic Dzy function, we crossed females that carried the *matGAL4* drivers, the *dzy* shRNA construct and that were heterozygous for a null allele of *dzy*, *dzy*^Δ*1*^, to males carrying the *dzy* shRNA construct and were also heterozygous for *dzy*^Δ*1*^ (Fig. S1). Thus all progeny would have strong maternal *dzy* RNAi knockdown and one quarter of progeny also would also be zygotically null for *dzy*, preventing any recovery of Dzy function. We assessed effectiveness of this by examining the cuticle phenotype, which allows us to assess effects on major morphogenetic movements and epithelial integrity. Wildtype embryos have an intact cuticle, secreted by the embryonic epidermis and have completed major morphogenetic movements like germband retraction, head involution and dorsal closure (Fig. 6K). Our previous cuticle analysis of *dzy*^Δ*1*^ maternal/zygotic null mutants revealed that head involution failed in essentially all embryos (Fig. 6L-N, yellow arrows; quantified in Fig. 6P), and in most embryos this was accompanied by small or large holes in the epidermis (Fig. 6M-N, red arrows), with a small fraction of embryos exhibiting more severe disruption of epidermal integrity (Fig. 6O). Intriguingly, this is less severe than full maternal/zygotic loss of Cno (Sawyer *et al*., 2009); instead it is more similar to the phenotype of CnoΔRA, which lacks the Ras-associated (RA) domains (Perez-Vale *et al*., 2021). Embryos from the *dzy^Δ1^ dzy* RNAi cross exhibited a similar range of phenotypes (Fig. 6P). The distribution of phenotypes was somewhat less severe in the progeny of the *dzy* RNAi cross than among *dzy*^Δ*1*^maternal/zygotic null mutants, but given that only 25% of he *dzy*^Δ*1*^ *dzy* RNAi embryos would be predicted to also be zygotically *dzy* null mutant, the overlap in cuticle phenotype suggests we have achieved strong reduction of maternal and zygotic Dzy function in this subset of embryos, and thus our strategy provided a reagent for defining the role of Dzy during morphogenesis. For all remaining experiments, we used the aforementioned *dzy*^Δ*1*^ *dzy* RNAi cross, referred to for simplicity below as “*dzy* RNAi embryos”.

### Like Cno, Dzy is required to reinforce junctions under tension during germband extension

Dzy is one of multiple Rap1 GEFs in the *Drosophila* genome. Our goal was to determine which of Cno’s many roles during morphogenesis require Dzy-mediated Rap1 activation and which may rely on other Rap1 GEFs. We envisioned three broad possibilities. First, given the multiple roles of Rap1 revealed above, if Dzy is the predominant GEF regulating Rap1 during morphogenesis, Dzy loss would lead to *Rap1*-like phenotypes and thus be more severe in its effects than Cno loss. Second, it was possible that Dzy activates Rap1 to mediate its action via Cno, but other GEFs regulate Rap1’s Cno independent roles. In this case Dzy loss would mimic loss of Cno. Finally, if Dzy was not an important Rap1 GEF during morphogenesis, *dzy* RNAi might not have later morphogenesis phenotypes.

We thus used the cross outlined above (Fig. S1) to strongly reduce maternal and zygotic Dzy and distinguish between these possibilities, beginning by exploring Dzy’s role during convergent elongation during germband extension, using our *dzy* RNAi approach. We began at the whole animal level. In *cno* mutants, germband elongation is significantly slowed (Sawyer *et al*., 2011). We thus assessed the effect of *dzy* RNAi on this process. To stage embryos, we used a “timer” that is likely to be independent of the cell movements we wanted to assess: the pattern of cell divisions in mitotic domains (Foe, 1989). We then measured the progress of the germband, normalizing it to the distance between the posterior end of the embryo and the cephalic furrow. While the degree of germband extension was similar between wildtype and *dzy* RNAi embryos at stage 7 (Fig. 7A-C), at both stages 8 (Fig. 7D-F) and 9 (Fig. 7G-I) germband extension in *dzy* RNAi embryos slowed substantially relative to wildtype. This delay resembles that seen after *cno* loss, which also preferentially affects the later stages of germband extension (Sawyer *et al*., 2011).

**Figure 7.**
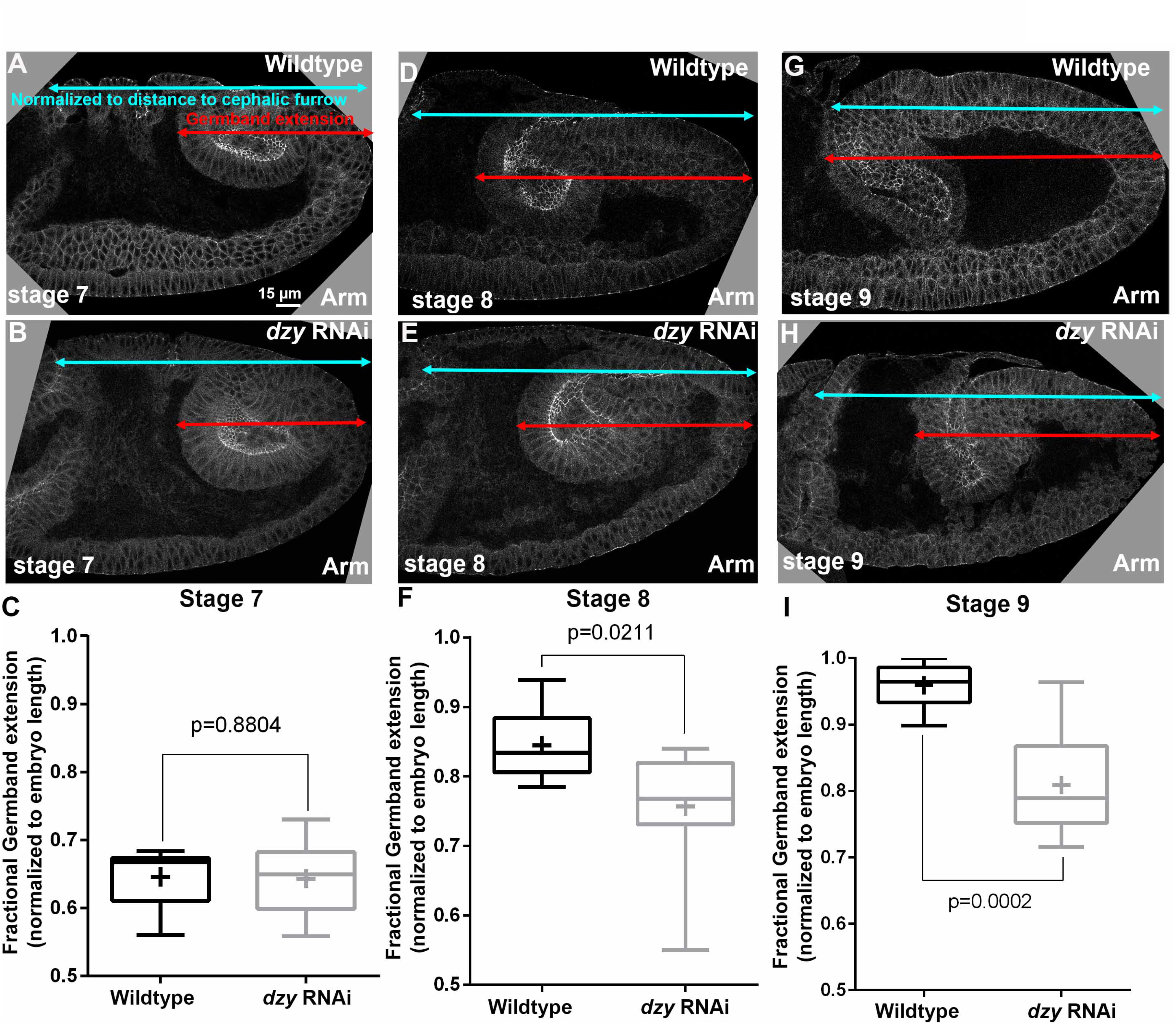
*dzy* RNAi delays germband extension. A, B, D, E, G, H. Cross sections of representative wildtype and *dzy* RNAi embryos, revealing extension of posterior end of the gut (red arrows) versus the full distance to the cephalic furrow (cyan arrows). C, F, I, Fractional germband extension at the indicated stages. *dzy* RNAi does not delay early germband extension (stage 7, C) but does reduce germband extension at later stages (F, I).

We next examined the effects of *dzy* RNAi on the orchestrated cell shape changes and AJ integrity during cell intercalation and germband extension. While loss of Dzy affects initial assembly of AJ and Cno proteins into apical AJs during cellularization, this is largely rescued as gastrulation begins (Bonello *et al*., 2018). Consistent with this, in *dzy* RNAi embryos the overall localization of both Arm (Fig. 6Q vs R, S; 8A’ vs B’) and Cno (Fig. 8A” vs B”) to AJs appeared relatively normal during gastrulation. As the germband extended in stage 7, we observed the first defects in *dzy* RNAi embryos: stacks of cells became abnormally elongated along the AP axis (Fig. 6R, S; 8B brackets). The dorsal folds also extended further ventrally than normal and ectopic folds were observed (Fig. 6R, S, 8B, red arrows; 14/21 stage 7 embryo had folds extending into the ectoderm). In wildtype embryos Arm and Cno were continuous around the cell circumference, both along aligned AP borders (Fig. 8A, red arrows) and at TCJs and short multicellular junctions (Fig. 8A, cyan arrowheads). In contrast, in *dzy* RNAi embryos gaps opened in apical AJs at many TCJs and AP borders (Fig. 8B, cyan arrowheads). All of these defects closely resembled phenotypes seen in *cno* mutants (Sawyer *et al*., 2011; Manning *et al*., 2019; Perez-Vale *et al*., 2021).

**Figure 8.**
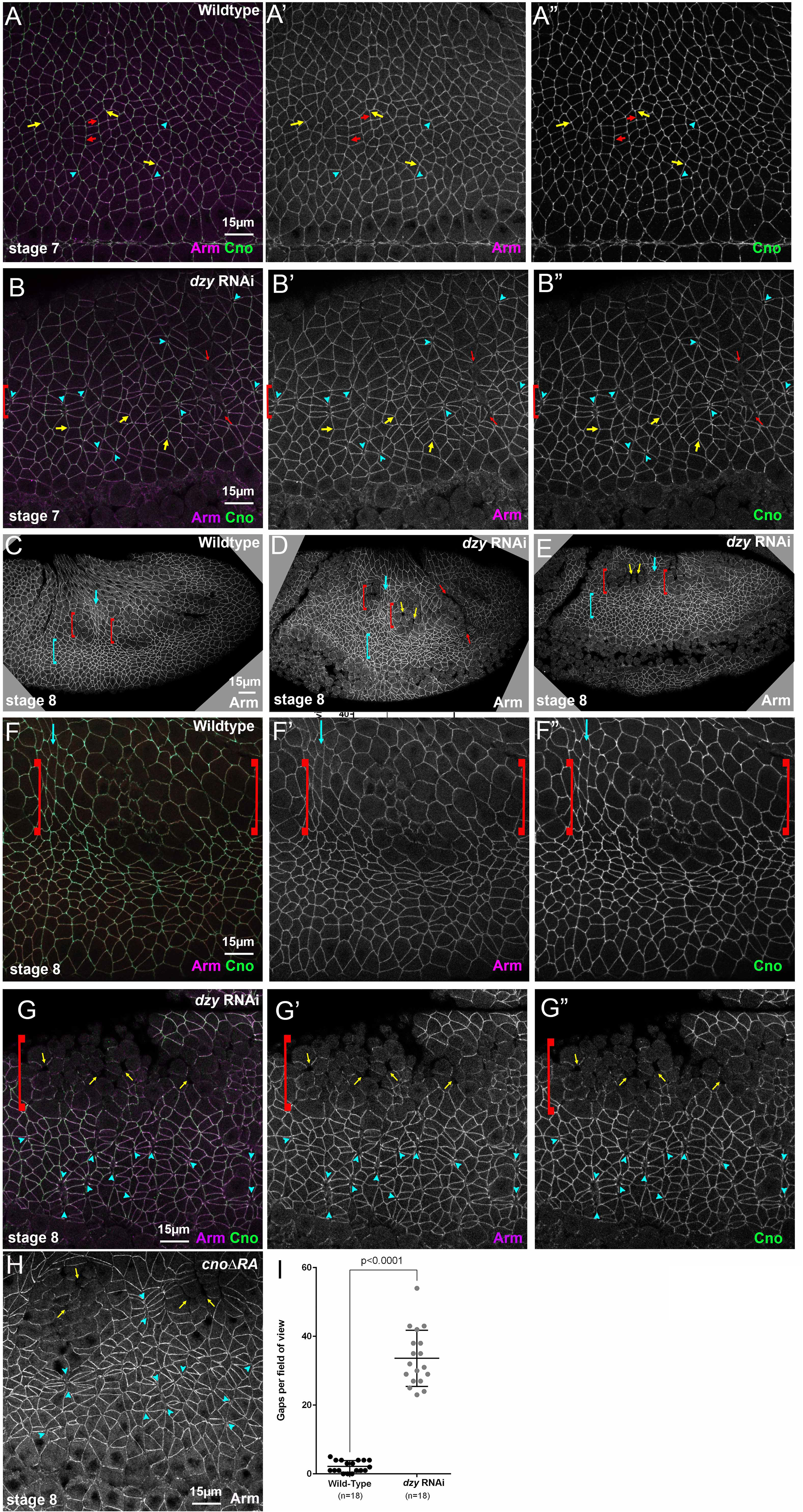
*dzy* RNAi leads to defects in cell shapes and at AJs under elevated tension similar to those seen in *cno* mutants. Embryos, stages 7 (A, B) and 8 (C-H). A. Wildtype closeup. Arm and Cno remain strong all around cells, including those at aligned AP borders (red arrows), those at TCJs (yellow arrows), where Cno is enriched, and at the centers of rosettes (cyan arrowheads). B. *dzy* RNAi. Gaps appear at aligned AP borders and rosette centers (cyan arrowheads), especially in regions where cells are abnormally elongated along the AP axis (brackets). Ectopic grooves are present (red arrows). Cno enrichment at TCJs is reduced (yellow arrows). C, F. At stage 8 in wildtype, cells in mitotic domain 11 round up to divide (red brackets) and cells between the domains apically constrict as their neighbors reduce contractility (cyan arrows). D, E, G. *dzy* RNAi. Note ectopic fold (red arrows), gaps between mitotic cells (yellow arrows), elongated cells in the ventral ectoderm (cyan brackets), and gaps at TCJs and aligned AP borders (cyan arrowheads). H. Similar gaps are seen in *cno*Δ*RA* mutants. I. Quantification of gaps.

In addition to the challenges of cell shape change and rearrangement, AJs also need to be remodeled as cells round up for cell division. During embryonic stage 8, cells in mitotic domain 11 round up to divide (Fig. 8C, F, red brackets), and intervening cells apically constrict (Fig. 8C, F, cyan arrows), perhaps due to less pulling force from mitotic neighbors (Ko *et al*., 2020). However, in wildtype embryos AJs remain intact around the circumference of mitotic cells, although protein localization per unit membrane is reduced (Fig. 8F, brackets). More ventral cells, which are yet to divide, continue intercalation and retain intact apical AJs throughout the process. In *dzy* RNAi embryos, mitotic domains divided on schedule (Fig. 8D, E, brackets), but gaps appeared between rounded up mitotic cells (Fig. 8D, E, J, yellow arrows). More ventrally, gaps remained in apical AJs at many TCJs and AP borders (Fig. 8G, arrowheads). Most embryos retained ectopic folds (Fig. 8D, red arrow; 11/15 stage 8 embryos). However, Arm and Cno continued to localize to AJs in places where gaps did not open (Fig. 8G). All of these defects were quite similar to those we previously observed in *cno*ΔRA embryos, lacking Cno’s Rap1-binding RA domains (Fig. 8H; (Perez-Vale *et al*., 2021). To quantitatively compare the effects on junctional stability of Cno and Dzy loss, we quantified gaps in wildtype versus *dzy* RNAi embryos (Fig. 8I). While wildtype embryos only had occasional gaps in apical junctions (2.2 gaps per 133 × 133 µm field of cells; n=18 stage 7 and 8 embryos), gaps were much more frequent in *dzy* RNAi embryos (33.6 gaps per 133 × 133 µm field of cells; n=18 stage 7 and 8 embryos). This was comparable to the average number of gaps we previously observed in both *cno*ΔRA embryos and *cno* maternal/zygotic null mutants (22 and 30 gaps per field, respectively; (Perez-Vale *et al*., 2021). Together these data suggest Dzy function is dispensable for Cno and Arm localization to AJs, but is important to reinforce AJs under elevated tension, like TCJs and constricting AP borders. This is consistent with Dzy being a major GEF involved in Rap1 regulation of Cno during these events. However, *dzy* RNAi embryos lacked the imbalanced apical contractility and fragmentation of junctional Baz seen after *Rap1* RNAi, and thus Dzy is clearly not the only GEF responsible for Rap1 activation during embryonic morphogenesis.

### Dzy is required for maintaining most but not all aspects of AJ planar polarity and Cno tension sensing

As noted above, planar polarization of AJ proteins and the actomyosin cytoskeleton are essential for cell intercalation during germband extension (Perez-Vale and Peifer, 2020). In wildtype embryos, Baz is enriched on DV borders (Fig. 9A, yellow vs cyan arrows: enrichment is ∼2-fold Fig. 9E), where it plays an important role in driving intercalation (Zallen and Wieschaus, 2004). Cno is important for maintaining balanced planar polarity. In Cno’s absence, E-cadherin (Ecad), Arm, and Pyd are reduced on AP borders, while Baz is essentially lost there, thus dramatically enhancing its planar polarization to DV borders (Sawyer *et al*., 2011; Perez-Vale *et al*., 2021). Given the parallel effects of loss of Cno and loss of Dzy on AJs under tension, we anticipated that effects on planar polarity of junctional proteins would be identical. However, when we measured junctional planar polarity, the results proved more complex. Like *cno* mutants, *dzy* RNAi embryos had significantly enhanced Baz planar polarity, with Baz strongly reduced or eliminated on AP borders (Fig. 9B, yellow vs cyan arrows; quantified in Fig. 9E). In wildtype, Pyd is also more subtly planar polarized, with enrichment on horizonal borders (Fig. 9C, yellow vs cyan arrows: Fig. 9F). Pyd planar polarity was also further enhanced by *dzy* RNAi (Fig. 9D, yellow vs cyan arrows; Fig. 9F), thus resembling the effect of *cno* mutants (Manning *et al*., 2019). However, in contrast to *cno* mutants (Sawyer *et al*., 2011), the planar polarity of Arm was not significantly enhanced in *dzy* RNAi embryos, when borders with obvious apical gaps were excluded (Fig. 9G).

**Figure 9.**
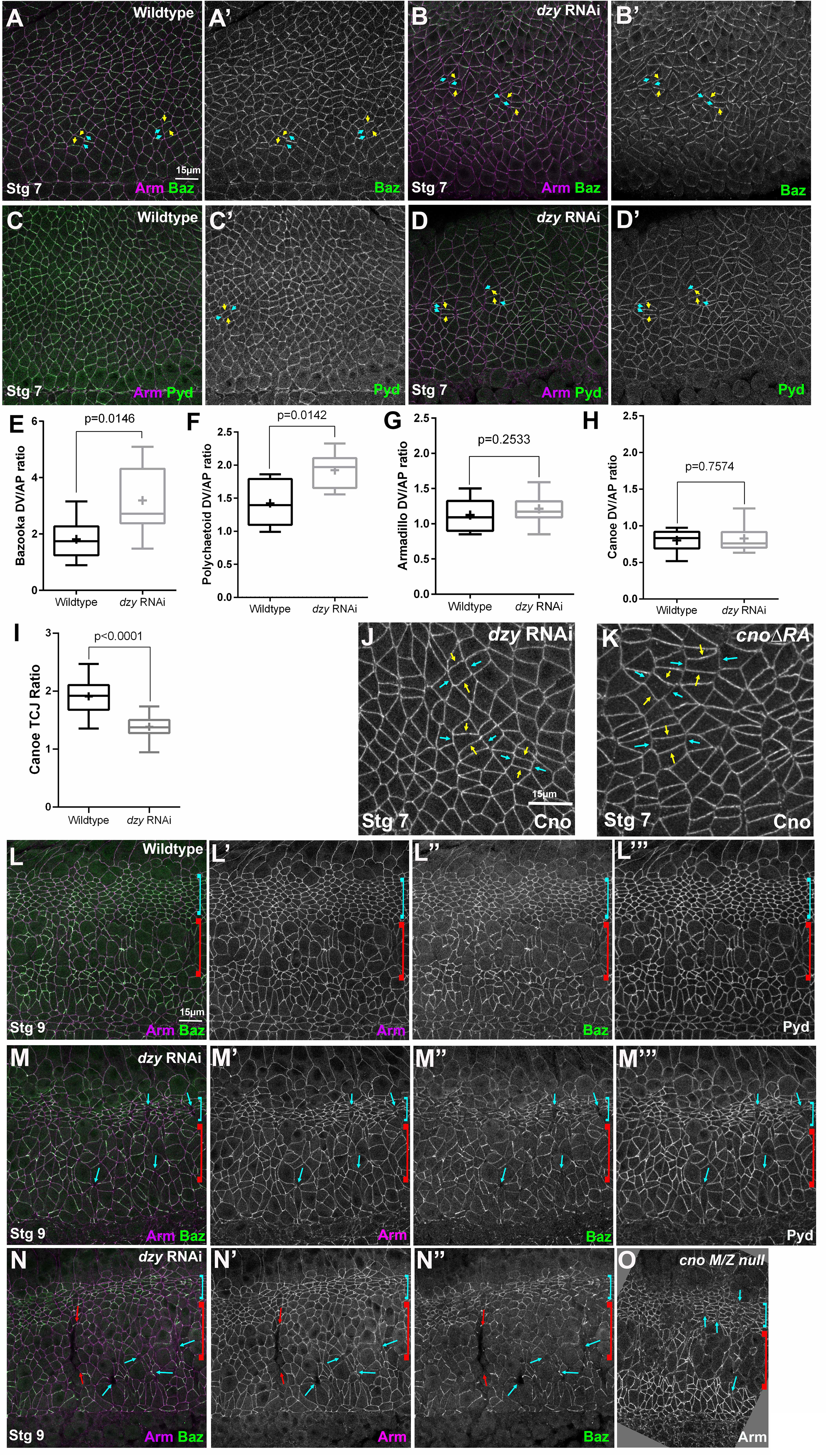
*dzy* RNAi alters junctional protein planar polarity in ways similar to but not identical to loss of Cno. A-D, J, K. Stage 7 embryos. A. Wildtype. Baz is planar polarized to DV borders (yellow vs cyan arrows). B. *dzy* RNAi. Baz planar polarization is enhanced. C. Wildtype. Pyd is mildly enhanced at DV borders (yellow vs cyan arrows). D. *dzy* RNAi. Pyd planar polarization is enhanced. E-H. Planar polarity quantification. I. *dzy* RNAi reduces Cno enrichment at TCJs. J, K. *dzy* RNAi does not alter Cno planar polarity (J), unlike *cno*Δ*RA* (K). L-N. Stage 9. L. Wildtype. Dorsal ectodermal cells in mitotic domain 11 resume columnar shape (cyan bracket), while neuroectodermal cells in mitotic domain N round up to divide (red bracket). M,N. *dzy* RNAi. Dorsal ectodermal cells are hyperconstricted (cyan brackets). Gaps (cyan arrows) and folds (red arrows) persist. O. *cno* M/Z null mutant for comparison. Numbers of cells analyzed for planar polarity are in Table S1.

Cno itself is not uniformly localized to AJs. Cno is enriched at TCJs (Sawyer *et al*., 2009) in a tension-sensitive manner (Yu and Zallen, 2020; Fig. 8A, yellow arrows; quantified in 9I), and also subtly planar polarized, with enrichment on AP borders in the wildtype, opposite to that of Baz (Sawyer *et al*., 2011; Fig. 9H). Both TCJs and AP borders are locations where tension is thought to be elevated (Fernandez-Gonzalez *et al*., 2009; Yu and Zallen, 2020; Perez-Vale *et al*., 2021). Cno enrichment at both TCJs and AP borders requires Cno’s Rap1-binding RA domain, as in *cno*Δ*RA* mutants Cno enrichment at TCJs is strongly reduced and planar polarity is flipped, as loss at AP borders means it is now enriched at DV borders (Perez-Vale *et al*., 2021; Fig. 9K yellow vs cyan arrows). Our data above confirm that this also requires Rap1. Based on this, we anticipated that Dzy would be required for both proper Cno TCJ enrichment and planar polarization. However, once again things were more complicated. Cno TCJ enrichment was strongly reduced after *dzy* RNAi (Fig. 8A vs B, yellow arrows; quantified in Fig. 9I). Intriguingly, however, Cno planar polarity was unchanged in *dzy* RNAi embryos (Fig. 9J; quantified in 9H), unlike what we observed in *cno*Δ*RA* mutants (Fig. 9J vs K). Together, these data suggest that regulation by Dzy is important for most but not all aspects of AJ protein localization, thereby paralleling the complexity we saw in the regulation of Cno by Rap1 versus Dzy during cellularization (Bonello *et al*., 2018).

### *dzy* RNAi embryos largely maintain epithelial integrity, with some defects ventrally

We finished this analysis by examining how loss of Dzy affected the maintenance of epithelial integrity as morphogenesis proceeded. Despite the defects in junctional integrity at AJs exposed to elevated tension in *dzy* RNAi embryos, the epidermal epithelium remained relatively intact at the end of germband extension (Fig. 8C vs D, E). In wildtype embryos at stages 9 and 10, dorsal ectodermal cells that divided in domain 11 have resumed columnar architecture (Fig. 9L, cyan bracket). During these stages, AJs are further challenged as cells in mitotic domain N (Fig. 9L, red bracket) and mitotic domain M round up to divide, and a subset of cells invaginate as neuroblasts. In wildtype embryos Arm and Pyd staining remain strong in both mitotic and non-mitotic cells while Baz is continuous in non-mitotic cells and reduced but still generally present at junctions of mitotic cells (Fig. 9L). In *dzy* RNAi embryos, while the ectoderm remained largely intact, multiple defects in cell shapes and AJ integrity persisted. Dorsal ectodermal cells were hyper-constricted along the AP axis (Fig. 9L vs M, N, cyan brackets), perhaps due to reduced pulling from mitotic neighbors. Gaps remained at TCJs and AP borders (Fig. 9M, N cyan arrows; Fig. 10B, arrowheads; 17/22 stage 9-10 embryos) and occasional ectopic grooves were present (Fig. 9N, red arrows; Fig. 10B arrow; 7/22 stage 9-10 embryos). Arm and Pyd staining remained largely continuous around cells, except at gaps (Fig. 9L, L”’ vs M, M”’; N”), and Baz staining, while less continuous (Fig. L” vs. M”,N”), was not fragmented as we observed in *Rap1* RNAi embryos. At this stage, defects were similar to those seen in *cno M/Z* null mutants (Fig. 9O).

**Figure 10.**
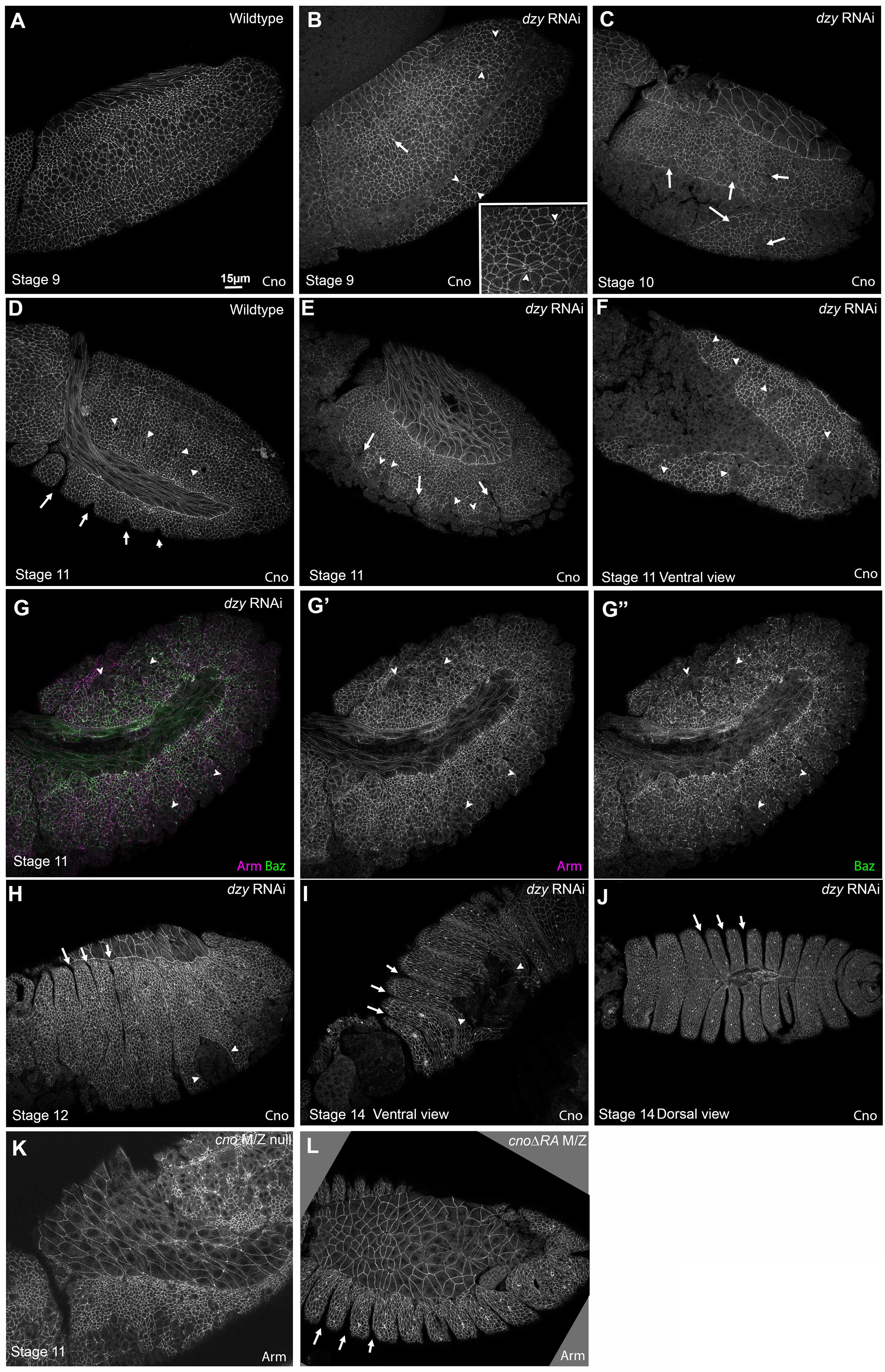
*dzy* RNAi leads to defects in the ventral epidermis and persistent deep segmental grooves. A, B. Stage 9. A. Wildtype. B. *dzy* RNAi. Note remaining fold (arrow) and gaps at some AP borders and TCJs (arrowheads, inset). C. *dzy* RNAi, stage 10. Gaps in the ventral epidermis are evident (arrows). D-G. Stage 11. D. Wildtype. Segment grooves become apparent (arrows) and tracheal pits invaginate (arrowheads). E, F, G. *dzy* RNAi. Note deep folds (arrows) and gaps in the ventral epidermis (arrowheads). H-J. *dzy* RNAi, stages 12 and 14. Note abnormally deep segmental grooves (arrows) and holes in the ventral epidermis (arrowheads). K. Ventral epidermal defects are more widespread in *cno M/Z* null mutants. L. *cno*Δ*RA* mutants share the deep segmental groove phenotype.

By stage 10, more severely affected embryos exhibited defects in epithelial integrity (Fig. 10C, arrows), perhaps where cells failed to regain columnar architecture following division. During stage 11, cell junctions in wildtype embryos are challenged by the formation of segmental grooves (Fig. 10D, arrows) and invagination of cells to form tracheal pits (Fig. 10D, arrowheads). At this stage, *dzy* RNAi embryo phenotypes become more variable, perhaps because RNAi knockdown is diminished in the 75% of embryos that are not *dzy* zygotic mutants. However, while epidermal integrity was broadly maintained, defects were observed in the ventral epidermis (Fig. 10E-G, arrowheads; 14/16 stage 11 and 12 embryos), often coinciding with the location of segmental grooves (Fig. 10E, arrows), which are also known to be sites of increased tension (Mulinari *et al*., 2008). These defects were less severe than those we observed in *cno^R2^* M/Z null mutants (Fig. 10K; Sawyer *et al*., 2009); instead they were more similar in severity to those observed in *cno*^Δ*RA*^ mutants (Fig. 10L; Manning *et al*., 2019; Perez-Vale *et al*., 2021). As in *cno* mutants (Manning *et al*., 2019), Baz localization to junctions was more affected than that of Arm (Fig. 10G), but we did not see the complete junctional fragmentation that we saw after Rap1 RNAi (above) or when we simultaneously reduced both Cno and Pyd (Manning *et al*., 2019). During dorsal closure, epidermal integrity remained largely intact, but many *dzy* RNAi embryos had ventral holes in the epidermis (Fig. 10H, I arrowheads), consistent with what we observed in the *dzy* RNAi cuticles. They also exhibited deep persistent segmental grooves (Fig. 10H, J arrows), another characteristic of *cno* mutants (Fig. 10L; Manning *et al*., 2019). However, consistent with their respective cuticle phenotypes, the effects of *dzy* RNAi on cuticle integrity and dorsal closure were less severe than those seen in *cno^R2^* M/Z null mutants (Fig 10E, G vs K).

Putting all of these observations together, the defects in morphogenesis seen in *dzy* RNAi embryos are similar to those of *cno* mutants, suggesting that Dzy is the major GEF acting via Rap1 to regulate Cno in these events. It’s intriguing that in many ways, especially the effect on overall epidermal integrity, *dzy* RNAi more closely resembles the phenotype of *cno*Δ*RA* rather than the even more severe effects of complete loss of Cno, suggesting Dzy may mediate Rap1 action via that direct interaction, with other GEFs regulating Rap1 and affecting Cno via less direct interactions. Thus, subtle differences in the effect on planar polarity of *dzy* RNAi and *cno*Δ*RA*, the reduced loss of epidermal integrity may suggest other GEFs act through Rap1 to regulate Cno. Finally, the substantially less severe phenotype of *dzy* RNAi relative to *Rap1* RNAi clearly suggest that Rap1 utilizes additional GEFS in regulating morphogenesis.

## Discussion

The diverse cell shape changes and arrangements during embryonic development require robust connections between cell-cell AJs and the contractile actomyosin cytoskeleton. AJs need to withstand the forces generated without disrupting tissue integrity. *Drosophila* Cno and its mammalian homolog Afadin are both essential for these events. One key challenge now is to define the mechanisms by which Cno is regulated. The small GTPase Rap1 regulates many events involving cell-cell and cell matrix adhesion, using diverse effectors, and Rap1, in turn, is activated by many different GEFs, acting at different times and places. Rap1 uses Cno as an effector during cellularization and mesoderm invagination, and during these stages Rap1 is regulated the PDZ-GEF Dzy. We sought to define Rap1’s roles in later morphogenesis, asking whether its roles match those of Cno, or whether it might play additional roles using other effectors. In parallel, we asked whether Dzy continued to be the predominant GEF involved in Rap1 regulation. Our results suggest that Rap1 has both Cno-dependent and Cno-independent roles in embryonic morphogenesis, allowing it to ensure tissue integrity. They further suggest Dzy is the main GEF involved in Cno regulation during these events, but that Rap1’s Cno-independent roles must involve another GEF.

### Rap1 regulates morphogenesis by restraining apical contractility, via Cno-dependent and Cno-independent effects

Given the parallel effects of loss of Cno and loss of Rap1 on cellularization and mesoderm apical constriction, we initially expected Rap1 loss to mimic loss of Cno. However, instead the phenotypes we observed suggest that Rap1 plays additional roles (Table 1). Some early effects of Rap1 loss parallel those of loss of Cno: gaps appear at many TCJs and AP borders, and AJ proteins and Baz planar polarity are altered in similar ways. However, quite rapidly additional defects appear in *Rap1* RNAi embryos that are not present in *cno* mutants. As gastrulation begins, apical cell shapes are drastically altered, with some cell apical areas much smaller and others much larger. This suggests unbalanced apical contractility. These differences are accentuated as subsets of cells round up to divide in mitotic domains, thus reducing their junctional contractility. Adjacent cells hyperconstrict, sometimes doing so to such an extent that groups of cells fold inward to form epithelial folds or balls. This may be due to reduced pulling forces across the tissue, both from the mitotic cells and from the open ventral furrow. It also could be that rounded up cells push on their neighbors. Many cells that rounded up to divide in stages 8 and 9 fail to restore columnar cell shape. As a result of these two defects, infolding of some cells and failure of others to resume columnar shape, tissue integrity is dramatically disrupted by stage 11. While loss of Cno or Dzy does lead to some unbalanced contractility and occasional tissue folds, these defects are much more severe and pervasive after Rap1 loss, providing strong evidence that Rap1 has additional effectors during morphogenesis.

**Table 1.** Comparison of the embryonic morphogenesis phenotypes of *cno M/Z* null mutants, *cno*Δ*RA* mutants, *Rap1* RNAi and *dzy* RNAi.

|  | ***cno* null**<br>(Sawyer et al. 2009; 2011;<br>Manning et al., 2019) | ***Rap1* RNAi**<br>(This work) | ***cno*Δ*RA***<br>(Perez-Vale et al. 2021) | ***dzy* RNAi**<br>(This work) |
| --- | --- | --- | --- | --- |
| Overall morphogenesis<br>as assessed by cuticles | Head involution and<br>dorsal closure failure<br>Moderate-Strong defects in<br>overall<br>epidermal integrity | Head involution and<br>dorsal closure failure<br>Moderate-Strong defects in<br>overall<br>epidermal integrity | Complete failure of both<br>head involution<br>and dorsal closure | Failure of head<br>involution. Dorsally closed<br>but frequent dorsal or<br>ventral holes |
| Cell shape defects and<br>gaps at AJs under<br>tension | Gaps common<br>(20-30 gaps/field of view)<br>but without widespread<br>unbalanced apical<br>constriction | Widespread unbalanced<br>apical constriction<br>and epithelial infolding<br>Gaps present in<br>less affected areas | Gaps common<br>(20-30 gaps/field of view)<br>but without widespread<br>unbalanced apical<br>constriction | Gaps common<br>(20-30 gaps/field of view)<br>but without widespread<br>unbalanced apical<br>constriction |
| Planar cell polarity<br>and AJ integrity |  |  |  |  |
| Baz | Planar polarity enhanced | AJ localization<br>strongly disrupted | Planar polarity enhanced | Planar polarity enhanced |
| Cno | NA | Planar polarity enhanced<br>and more punctate | Planar polarity enhanced | Planar polarity unaffected |
| Arm | Planar polarity enhanced | Planar polarity enhanced<br>and more punctate | Planar polarity enhanced | Planar polarity unaffected |
| Epidermal<br>integrity | Frequent large holes<br>in the ventral epidermis | Only dorsal epidermis<br>remains intact | Small holes<br>in the ventral epidermis | Small holes<br>in the ventral epidermis |

Rap1’s effects on AJ protein localization provide some clues. Cno loss accentuates the planar polarity of AJ proteins and Baz, but remain localized to AJs. In contrast, after *Rap1* RNAi Baz localization is affected much earlier and much more dramatically. While Baz returned to apical junctions after the end of cellularization after *Rap1* RNAi (Bonello *et al*., 2018), as germband extension began, Baz localization in *Rap1* RNAi embryos rapidly became discontinuous. This fragmentation persisted and was accentuated by stages 8 and 9. In contrast, Arm, Cno and Pyd remain localized to junctions, though their accumulation around the cell circumference is less regular than is true in wildtype—all are enriched at places where Baz remains. This suggests that effects of Rap1 loss on Baz are not simply the consequence of alterations in AJs, but, instead, that Rap1 may have more direct effects on Baz. However, Rap1 loss does not fully replicate the effects of total maternal/zygotic Baz loss, which leads to complete fragmentation of the epithelium and the cuticle it secretes (Müller and Wieschaus, 1996; Harris and Peifer, 2004). One speculative possibility is that the initial apical re-localization of Baz to AJs allows their assembly, but after Baz fragmentation, AJs lose the ability to balance contractility among neighbors. Consistent with Rap1 and Baz operating together or in parallel in balanced contractility, both Rap1 and Baz loss lead to increased variability of apical cell shape at cellularization (Choi *et al*., 2013). Rap1 has also been demonstrated to regulate AJ accumulation of both Arm and Baz in the developing adult photoreceptors (Walther *et al*., 2018). However, in that tissue Cno appears to be the relevant Rap1 effector involved in regulating Baz.

One important task going forward will be to identify additional Rap1 effectors. At this point, we can only speculate about possibilities. Combined reduction of Cno and loss of the ZO-1 homolog Pyd also leads to dramatic defects in epithelial integrity and infolding of epidermal cells, effects that are much more severe than loss/reduction of either alone (Manning *et al*., 2019) but are more similar to the effects of Rap1 loss. Could Pyd be another Rap1 effector in the *Drosophila* ectoderm? In mammalian cultured epithelial cells and in the developing mouse lens epithelium, Rap1 can regulate ZO-1 localization (Maddala *et al*., 2015). However, while combined reduction of Cno and loss of Pyd disrupts Baz in a similar way to loss of Rap1, its effect on Arm localization is more dramatic—both Arm and Baz localization to AJs are fragmented after combined reduction of Cno and loss of Pyd (Manning *et al*., 2019). Another speculative possibility is that aPKC might be a Rap1 effector. In cultured mammalian cells, Rap1 can act through the adapter Shank, which binds aPKC and regulates junctional integrity (Sasaki *et al*., 2020). However, complete loss of aPKC has earlier and more severe consequences in ectodermal integrity in *Drosophila*, with complete loss of epithelial integrity by stage 9 (Harris and Peifer, 2007). One intriguing possibility, inspired by a recent preprint from the Morais-de-Sá lab, is that loss of Rap1 reduces but does not eliminate aPKC function (https://www.biorxiv.org/content/10.1101/2022.03.02.482459v1). They examined a different *Drosophila* epithelium—the ovarian follicle cells. Reducing aPKC activity led to phenotypes strikingly reminiscent to those we saw in the embryo after Rap1 RNAi—epithelial contractility became unbalanced, with cells next to mitotic cells undergoing unrestrained apical constriction. Another way Rap1 could regulate apical contractility is via effects on Rho, the activity of which it is known to regulate in endothelial cells (Pannekoek *et al*., 2014). It will be of interest to explore these and other speculative possibilities.

### Dzy is the predominant GEF involved in Cno regulation during morphogenesis, but Rap1 must use additional GEFs during these events

Rap1 has many known GEFs that – in both flies and mammals – play roles at different times and in different tissues. Defining the relevant GEFs that regulate junction-cytoskeletal linkage during embryonic morphogenesis remains an important task. Previous work examined the two earliest stages of *Drosophila* development, cellularization and mesoderm invagination, and the answer was complex. During cellularization, we know of two GEFs that act sequentially: the atypical GEF Sponge regulates initial placement of Cno and AJ proteins prior to membrane invagination (Schmidt *et al*., 2018), while Dzy regulates correct assembly of nascent AJs (Bonello *et al*., 2018). However, loss of Dzy does not affect either Cno localization or AJs placement as dramatically as loss of Rap1. Another complexity emerges from the fact that Rap1 is essential for Cno recruitment to the plasma membrane, while Cno remains localized to the membrane in *dzy* mutants, but its precise apical positioning and enrichment at TCJs is disrupted. The effect of Dzy loss on AJ protein localization is also somewhat milder than the effect of loss of Rap1. Intriguingly, the phenotype of Dzy loss mimics that of the loss of Cno’s Rap1-binding RA domains. These data suggested that during cellularization Rap1 affects Cno and AJ positioning via both RA-dependent and RA-independent mechanisms, and that Dzy mediates the RA-domain-dependent effects, while other GEFs act in parallel to regulate Rap1 and mediate its RA-domain-independent regulation of Cno, likely via other intermediate effectors. In contrast, loss of Dzy, Cno and Rap1 have similar effects on mesoderm invagination (Sawyer *et al*., 2009; Spahn *et al*., 2012).

Our current work suggests an even greater level of complexity of Rap1 regulation and action during germband extension and the maintenance of epithelial integrity. Loss of Dzy has effects similar to those of loss of Cno, in terms of altered cell rearrangements, stability of AJs under elevated tension, and planar polarization of junctional proteins (Table 1). However, some effects of Dzy loss do not fully replicate the effects of complete loss of Cno. These include a lack of effect on Arm planar polarity, and differences in the degree of loss of epithelial integrity. While it is possible some of these differences are due to a failure to completely eliminate Dzy using our knockdown/zygotic mutant approach, we think it is more likely that Cno retains some residual activity in Dzy’s absence. Intriguingly, the somewhat less severe effects of *dzy* RNAi on epithelial integrity are shared by *cno*Δ*RA* mutants. Thus, Dzy may be required for the “direct” Cno activation mediated by active Rap1 bound to the RA-domains, but Rap1 may also activate Cno to a lesser extent via RA-domain and Dzy-independent mechanisms, presumably when it is activated by a different GEF. This latter regulation may be indirect, via the other speculative effectors discussed above. These data also raise another intriguing possibility: that different pools of Rap1 at different subcellular locations carry our Rap1’s Cno-dependent and Cno-independent roles, and that different GEFs localize to and act at these different locations. In future studies, it will be important to identify those GEFs, and, as we discussed above, to identify Rap1’s additional effectors involved in regulating morphogenesis.

## Materials and methods

### Fly stocks and generation of the *dzy* shRNA construct

We generated an UAS-driven 21 nucleotide siRNA targeting a region of the 5^th^ exon of dizzy. We used the Designer of Small Interfering RNA (DSIR: http://biodev.extra.cea.fr/DSIR/DSIR.html) tool to design the 21 nucleotide target. The target sequence was input into the Predicted Off-Target Free Sequence Regions (https://www.flyrnai.org/RNAi_find_frag_free.html). The targeted sequence is **cgttctatccgatcgtcaaga** (see additional oligo design information below). We followed the TRiP protocols (Perkins *et al*., 2015), cloning the hairpins into the WALIUM20 vector. The UAS driven siRNA was inserted by BestGene Inc. into the 3rd chromosome attP docking site for phiC31 integrase-mediated transformation (position 68A4; Bloomington Drosophila Stock Center stock #8622 (genotype: *y^1^ w^67c23^; P{CaryP}attP2*).

Oligo Design is done as follows:

Top strand oligo: ctagcagt—**insert sense oligo**—tagttatattcaagcata—**insert anti-sense oligo**—gcg
Bottom strand oligo: aattcgc—**sense oligo**—tatgcttgaatataacta—**anti-sense oligo**—actg
For targeting *dzy*, this involved the following oligonucleotides:
Top strand oligo: ctagcagt**cgttctatccgatcgtcaaga**tagttatattcaagcata**tcttgacgatcggatagaacg**gcg Bottom strand oligo: aattcgc**cgttctatccgatcgtcaaga**tatgcttgaatataacta**tcttgacgatcggatagaacg**actg

We used *yellow white* flies as our control. These are referred to as Wildtype (WT) in figures and text. All experiments involving *dzy* RNAi and the associated WT embryos were conducted at 27°C, to enhance GAL4 activity. The other experiments were performed at 25°C. The fly stocks used and their sources are outlined in Table S2. *Rap1* knockdown by shRNA was completed by crossing double-copy *mat*-*tub*-*GAL4* females to UAS.Rap1 RNAi v20/TM3, Sb males (Figure S1).

### Embryo fixation and immunofluorescence

Flies were crossed in cages over apple juice agar plates with yeast paste and left to lay eggs for 4-18 hours before collection. Our method of embryo collection, embryo fixation, and embryo staining was previously described by (Bonello *et al*., 2018). Briefly for heat-fixation: We removed the chorion membrane by nutating in 50% bleach. Afterwards, the embryos were washed three times in 0.03% Triton X-100 with 68 mM sodium chloride (NaCl), and then fixed in 95°C Triton salt solution (0.03% Triton X-100 with 68 mM NaCl and 8 mM EGTA) for 10 seconds. We fast-cooled samples by adding ice-cold Triton salt solution and placing on ice for at least 30 minutes. We removed the vitelline membrane by vigorously shaking in 1:1 heptane/methanol solution. The embryos were again washed thrice with 95% methanol/5% EGTA and stored in 95% methanol/5% EGTA for at least 24 hours at −20°C before staining. Before staining, the heat-fixed embryos were washed three times with 0.01% Triton X-100 in PBS (PBS-T). For embryos that were formaldehyde-fixed (*Rap1* RNAi when compared with His::RFP): embryos were dechorionated in 50% bleach, washed three times in 0.1% Triton X-100, and then fixed in 8% formaldehyde, 1X PBS, and 8 mM EGTA for 30 minutes. These embryos were devitellinized by vigorous shaking in 1:1 heptane/90% ethanol. The embryos were finally washed three times with 90% ethanol and stored in 90% ethanol for at least 24 hours at −20°C before staining. At the point of staining, the formaldehyde fixed embryos were washed three times with 0.1% Triton X-100 in PBS. All embryos were blocked in 1% normal goat serum in PBS-T (PNT) for 1 hour. The embryos were then nutated in the primary antibodies overnight at 4°C, washed with PNT, and nutated in secondary antibodies overnight at 4°C. We used PNT to dilute the primary and secondary antibodies. After incubating in the secondary antibody, the embryos were washed with PNT and stored in 50% glycerol at −20°C. Later, we used Aquapolymount (Polysciences) to mount onto glass slides. The antibodies used and their sources are outlined in Table S3.

### Image acquisition and analysis

All images were obtained from fixed embryos. We imaged on the Carl Zeiss LSM 880 confocal laser-scanning microscope. Images were captured on the 40x/1.3 NA Plan-Apochromat oil objective. Brightness and contrast were fine-tuned using both ZEN 2009 software and after image processing in Adobe Photoshop. To capture the enrichment of proteins at the adherens junctions, we created maximum intensity projections (MIPs) with tools from ImageJ (National Institutes of Health). MIPs for Cno TCJ enrichment and planar-cell-polarity analysis were created from Z-stacks (1024 x 1024 pixels) through the ventrolateral ectoderm of the embryo at stages 7 and 8 respectively.

### Cno TCJ enrichment analysis

We obtained TCJ enrichment data from MIPs, using a digital zoom of 1.6 and a step size of 0.3 µm. First, we determined the length of the AJs through the embryo, which was on average 7 slices from the most apical portion of the junction to the most basal, spanning 1.5-2.4 µm of the apical region of stage 7 epithelial cells. We captured the mean intensity of Cno in ImageJ software by measuring the pixel intensity along the lines of the cell borders using the line tool at a width of 3 pixels. We next created a short line at the TCJ, careful to avoid overlapping the bicellular border lines. For each TCJ or multicellular junction we analyzed, the connecting three or four bicellular junctions were measured for comparison. We quantified 10 cells per embryo measured, and a total of four (*dzy* RNAi) or five (*Rap1* RNAi) embryos were assessed from three independent experiments. We calculated the Cno TCJ ratio by dividing the mean pixel intensity of the TCJ by the average intensity of the bicellular junctions. Statistical analysis was calculated by GraphPad Prism 9. Significance was determined by Welch’s corrected unpaired *t* test. We did not assume equal SD. We generated box and whisker plots using GraphPad Prism software, where the box represents the 25^th^ to 75^th^ percentile, the whiskers represent 5^th^ to 95^th^ percentiles, the median is displayed by a horizontal line within the box, and the mean is represented by a plus sign (+).

### Gap analysis

Data for gaps per field of view was collected visually at the level of apical AJs. A field of view measured 133 x 133 µm, collected from Z-stacks at the ventrolateral epidermis of each embryo. Data was collected from embryos at stages 7 and 8. Gaps were visualized using the Arm channel. Gaps formed at TCJs, rosette centers, or along AP borders spanning several cells (exceeding ∼1 µm). Along AP borders, gaps that transversed the length of about 4 cells (likely the location of a failed rosette) were considered one gap. Graphs and statistical analysis were executed in GraphPad Prism software, using Welch’s unpaired *t* test.

### Germband extension analysis

Cross section and apical view images of embryos were used to determine the rate of germband extension as compared to embryo stage. Three timepoints of germband extension were chosen: stage 7, 8, and 9. These stages were matched across mutant and WT embryos using the mitotic domains as described by VE Foe (1989). The length of the embryo was normalized as the distance between the posterior end and the cephalic furrow on the dorsal side of the embryo. The degree of germband extension at stage 7, 8, or 9 was quantified as the percent of the length of the embryo that the anterior-most columnar gut cells had reached at the time of fixation. This was measured by creating vertical lines in ImageJ at the posterior end of the embryo, at the cephalic fold, and at the anterior-most gut cells. We then created perpendicular lines to measure the pixel distance between each specified point for the length of the embryo and the length of germband extension. The graph was plotted as average length of germband extension divided by average normalized length of embryo. Data statistical analysis was done using GraphPad Prism 9. Statistical significance was calculated by Welch’s unpaired *t* test.

### Planar polarity quantification

We collected data for planar polarity from MIPs of a 2.4 µm apical section of AJs from embryos at stage 8. The MIPs were generated with the Stacks tool in ImageJ after determining the length of the AJs from Z-stacks through the embryo. Then we used the line tool at a width of 3 pixels to measure the mean intensity of protein fluorescence at both the AP and DV borders of a selected cell, careful to avoid overlapping the TCJ/multicellular junctions. We distinguished each border by their angle. Borders angled from 60°–90° were considered vertical AP borders, while borders angled at 0°–29° were consider horizontal DV borders. We then used the Circle tool in the cytoplasmic region of 13 cells to gather the mean background intensity. We subtracted the background intensity for each antigen from the intensity at the bicellular borders to get our results for Cno, Baz, Arm, and Pyd planar polarity. For *dzy* RNAi, a total of 21 embryos were assessed from at least 6 different experiments. For *Rap1* RNAi, a total 12 embryos were assessed from at least 7 different experiments. The number of cells assessed per antigen can be found in Table S1. We normalized our proteins of interest to the AP borders and expressed our results as a DV/AP ratio. We generated box and whisker plots to present statistical data using GraphPad Prism software, where the box represents the 25^th^ to 75^th^ percentile, the whiskers represent 5^th^ to 95^th^ percentiles, the median is displayed by a horizontal line within the box, and the mean is represented by a plus sign (+).

### Cuticle preparation

Cuticle preparation was performed according to (Wieschaus and Nüsslein-Volhard, 1986). In brief, embryos were aligned on agar plates and allowed to develop fully. Unhatched embryos were nutated in 50% bleach to remove the chorion membrane. After nutating, the embryos were washed in 0.1% Triton X-100 and mounted on glass slides in 1:1 Hoyer’s medium/lactic acid. The glass slides were incubated at 60°C for 48 hours, then stored at room temperature.

## Acknowledgements

We are grateful to Peifer lab members for helpful advice and discussions, to Rodrigo Fernandez-Gonzalez, Katheryn Rothenberg, Kevin Slep for important feedback on the manuscript, and the Bloomington Drosophila Stock Center for fly stocks. We thank Tony Perdue of the Biology Imaging Center for confocal imaging advice and support. This work was supported by NIH R35 GM118096 to M.P. K.Z.P.-V. was supported by the National Institutes of Health F31 GM131521 and by a Graduate Diversity Enrichment Program Award from the Burroughs Wellcome Fund. The authors declare no competing financial interests.

## Author contributions

K.Z. Perez-Vale and M. Peifer conceived the study, K.Z. Perez-Vale designed the Dzy knockdown strategy, carried out the initial experiments with assistance from M. Greene and N. J. Gurley, and trained K.D. Yow. K.D. Yow carried out most of the analysis of the *Rap1* and *dzy* phenotypes. K.Z. Perez-Vale, K.D. Yow, and M. Peifer drafted the manuscript with input from the other authors.

---

**Figure S1.**
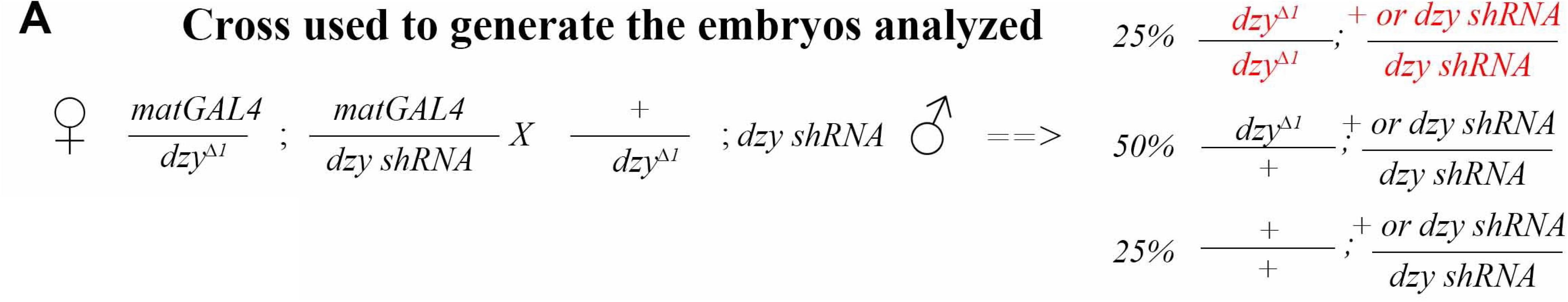
Genetic cross used to generate “dzy RNAi” embryos. E used the GAL4/UAS system to express an shRNA targeting dzy both maternally and zygotically. The use of the strong dual MatGAL4 driver ensures strong knockdown of maternal mRNA. 25% of the embryo are homozygous for the *dzy*^Δ*1*^ null allele and thus have no zygotic gene expression—these should have the strongest knockdown of Dzy protein.

**Table S1.** Number of cells analyzed for planar cell polarity.

| **<u>Antigen</u>** | **<u>Wildtype</u>** | **<u>*dzy* RNAi</u>** | **<u>*rap1* RNAi</u>** |
| --- | --- | --- | --- |
| Armadillo | 129 | 236 | 145 |
| Bazooka | 65 | 111 | 72 |
| Canoe | 64 | 125 | 73 |
| Polychaetoid | 64 | 125 | 73 |

**Table S2.** *Drosophila* stocks used in this study.

| *Drosophila* Stock | Source |
| --- | --- |
| *dzy*<sup>Δ1</sup>/*Cyo* | <u>Huelsmann et al., 2006</u> |
| *dzy shRNA* | Created here—see Materials and Methods |
| *y*<sup>1</sup> *w*<sup>67c23</sup>; *P{CaryP}attP2* | Bloomington Drosophila Stock Center (Stock #8622) |
| *Mat-tub-Gal4;Mat-tub-Gal4* | Bloomington Drosophila Stock Center (stock #80361) |
| *UAS.Rap1 RNAi v20/TM3, Sb* | Bloomington Drosophila Stock Center (stock no. 35047) |

**Table S3.** Antibodies used in this study.

| Primaries | Species | Dilution | Source |
| --- | --- | --- | --- |
| Anti-Canoe | Rabbit | 1:1,000 | Sawyer et al., 2009 |
| Anti-Bazooka | Rabbit | 1:2,000 | Choi et al., 2013 |
| Anti-Armadillo<br>(N27A1) | Mouse2a | 1:100 | Developmental Studies Hybridoma Bank |
| Anti-Polychaetoid<br>(PYD2) | Mouse2b | 1:500 | Developmental Studies Hybridoma Bank |
| Secondary antibodies |  | Dilution | Source |
| Alexa Fluor 488, 568, 647 |  | 1:1,000 | Life Technologies |

